# Repeatability of adaptive radiation depends on spatial scale: regional versus global replicates of stickleback in lake versus stream habitats

**DOI:** 10.1101/564005

**Authors:** Antoine Paccard, Dieta Hanson, Yoel E. Stuart, Frank A. von Hippel, Martin Kalbe, Tom Klepaker, Skúli Skúlason, Bjarni K. Kristjánsson, Daniel I. Bolnick, Andrew P. Hendry, Rowan D.H. Barrett

## Abstract

The repeatability of adaptive radiation is expected to be scale dependent, with determinism decreasing as greater spatial separation among “replicates” leads to their increased genetic and ecological independence. Threespine stickleback (Gasterosteus aculeatus) provide an opportunity to test whether this expectation holds for the early stages of adaptive radiation -their diversification in freshwater ecosystems has been replicated many times. To better understand the repeatability of that adaptive radiation, we examined the influence of geographic scale on levels of parallel evolution by quantifying phenotypic and genetic divergence between lake and stream stickleback pairs sampled at regional (Vancouver Island) and global (North America and Europe) scales. We measured phenotypes known to show lake-stream divergence and used reduced representation genome-wide sequencing to estimate genetic divergence. We assessed the scale-dependence of parallel evolution by comparing effect sizes from multivariate models and also the direction and magnitude of lake-stream divergence vectors. At the phenotypic level, parallelism was greater at the regional than the global scale. At the genetic level, putative selected loci showed greater lake-stream parallelism at the regional than the global scale. Generally, the level of parallel evolution was low at both scales, except for some key univariate traits. Divergence vectors were often orthogonal, highlighting possible ecological and genetic constraints on parallel evolution at both scales. Overall, our results confirm that the repeatability of adaptive radiation decreases at increasing spatial scales. We suggest that greater environmental heterogeneity at larger scales imposes different selection regimes, thus generating lower repeatability of adaptive radiation at larger spatial scales.

## INTRODUCTION

An unanswered question surrounding adaptive radiation is the extent to which the process is deterministic versus stochastic. That is, if the “tape of life” were rewound and started over multiple independent times, to what extent would a similar set of species evolve when presented with a similar set of environmental niches (Gould 1990; Blount *et al.*, 2018)? This thought experiment cannot be made real outside of the laboratory (MacLean, 2005), and so evolutionary biologists have tended to substitute geographical space for evolutionary time by asking: To what extent do related species evolve in parallel in similar sets of environments that are spatially isolated and evolutionarily independent? A difficulty in addressing this question is that relatively few adaptive radiations are replicated in space; many are instead unique, such as Darwin’s finches in the Galapagos Islands (Grant 1981) and the Hawaii honeycreepers (James 2004); or are relatively rare, such as the radiations of cichlids in the three Great Lakes of Africa (Kocher 2004). A few adaptive radiations are sufficiently replicated to make inferences about repeatability; and these cases generally indicate that adaptive radiations have substantial repeatable components, such as the same ecomorphs of *Anolis* lizards evolving on multiple islands in the Greater Antilles (Losos 2009) or the same ecomorphs of spiders evolving on multiple Hawaiian Islands (Gillespie, 2004; Gillespie *et al.*, 2018). Yet these same radiations often show substantial non-deterministic components, such as “missing” ecomorphs of lizards or spiders on some islands or strong contingency depending on physical properties of the system (Brawand *et al.* 2014).

A remaining complication is that repeated adaptive radiations with a given group of organisms are not entirely independent – genetically or ecologically. With respect to genetic non-independence, post-colonization gene flow occurs among islands for *Anolis* lizards (Losos 2009) and Hawaiian spiders (Gillespie 2004), and independent radiations of African cichlids appear to be using some of the same ancestral polymorphisms (Loh *et al.* 2012). With respect to ecological non-independence, adaptive radiations are often replicated on only regional spatial scales (e.g., among islands in Caribbean or Hawaii), where overall ecological conditions are relatively similar. Hence, the resulting genetic or ecological non-independence could be primary factors shaping deterministic responses. Stated in the same counterfactual sense as “replaying the tape of life”, if a few ancestral species had colonized very different geographic locations, to what extent would a similar adaptive radiation take place? Intuition would suggest that such “replicate” adaptive radiations separated by large geographical scale would yield lower repeatability than would replicate adaptive radiations at a more regional scale – as a result of greater genetic and ecological independence.

The multi-scale adaptive radiation thought experiment described above is not inevitably counterfactual. Threespine stickleback (*Gasterosteus aculeatus*) are found throughout much of the northern hemisphere, where they show many replicate adaptive radiations in freshwater ecosystems arising from independent colonization events from a common marine ancestor (Colosimo *et al.* 2005). Although the resulting populations are all placed under the same Latin binomial, many of these populations show large genetic differences that are primarily shaped by adaptation to different environments as opposed to interactions between close relatives, as in the Hawaiian spiders or Anolis lizards (Gillespie, 2004; Losos, 2009; Schluter *et al.*, 2010; Peichel & Marques, 2017; Gillespie *et al.*, 2018). For example, the F_ST_ between parapatric lake-stream stickleback is greater than 0.10 in each of many watersheds colonized postglacially by marine stickleback (Kaeuffer *et al.*, 2012; Roesti *et al.*, 2012; Stuart *et al.*, 2017). Hence, different freshwater populations of stickleback can be thought of as representing the early stages of adaptive radiation (Schluter & McPhail 1992; Schluter 1996; Taylor & McPhail 1999). With this stickleback system, one can compare regional replicate adaptive radiations (stickleback in similar habitats in different watersheds in the same geographical area) with global replicate adaptive radiations (stickleback in similar habitats in different watersheds in distant geographical areas). We therefore use this system to ask whether genetic and phenotypic divergence is more repeatable (similar magnitude and direction across replicates) at regional than at global scales.

### Study system

Parapatric pairs of lake and stream stickleback have been used frequently to study the repeatability/predictability/parallelism/convergence of adaptive divergence (e.g., Reimchen *et al.* 1985; Lavin & McPhail 1993; Deagle *et al.* 1996; Thompson *et al.* 1997; Reusch *et al.* 2001; Hendry & Taylor 2004; Aguirre 2009; Lucek *et al.* 2010; Eizaguirre *et al.* 2011; Kaeuffer *et al.* 2012; Roesti *et al.* 2012; Ravinet *et al.* 2013; Lucek *et al.* 2013; Weber *et al.* 2017; Stuart *et al.* 2017). In this study system, stickleback in a lake and its adjoining inlet or outlet stream (hereafter, lake-stream population pairs) often evolve similar patterns of phenotypic divergence in independent watersheds. Most notably, lake stickleback generally evolve shallower bodies and more gill rakers than do stream stickleback, a pattern driven at least in part by consistent differences in diet between habitats (Berner *et al.* 2008; Kaeuffer *et al.* 2012). These phenotypic differences tend to be genetically based (Sharpe *et al.* 2008; Hendry *et al.* 2011; Berner *et al.* 2011; Oke *et al.* 2016; Moser *et al.* 2016) and are likely adaptive, given their close association with variation in prey availability (Berner *et al.* 2008; Kaeuffer *et al.* 2012). However, non-parallel aspects of divergence are known where some traits show low levels of parallelism (Stuart *et al.* 2017).

Most analyses of parallel evolution in lake-stream stickleback have taken place on regional geographic scales, where we expect generally similar selective regimes and genetic backgrounds: northern Vancouver Island (Lavin & McPhail 1993; Hendry & Taylor 2004; Berner *et al.* 2008; 2009; Kaeuffer *et al.* 2012; Stuart *et al.* 2017), Haida Gwaii (Reimchen *et al.* 1985), Ireland (Ravinet *et al.* 2013), and northern Germany (Reusch *et al.* 2001). Several studies, however, have analyzed sets of lake-stream pairs within and between regions: Vancouver Island vs. Switzerland (Berner *et al.* 2010), Vancouver Island vs. artificial habitats in California (Hendry *et al.* 2013), Iceland vs. Switzerland (Lucek *et al.* 2014b), and Vancouver Island vs. Switzerland vs. Ireland (Lucek *et al.* 2013). These studies revealed important morphological and genetic variation in the direction and magnitude of lake-stream divergence between regions, but they did not formally compare the extent of parallel evolution across different geographic scales. In the present study, we conduct such a comparison by assessing parallel evolution of lake-stream divergence at two very different scales: six pairs within Vancouver Island (hereafter, “regional” comparisons) and six pairs from worldwide localities that encompass the majority of the threespine stickleback range (hereafter, “global” comparisons).

We estimated phenotypic (non)parallelism by first comparing the effect sizes of multiple univariate and multivariate models. We then evaluated the degree of phenotypic and genetic (non)parallelism by calculating multivariate phenotypic and genetic (neutral and outlier) lake-to-stream divergence vectors. Then, we quantified the differences among vectors in direction (i.e., the angle between any two divergence vectors) and in magnitude (i.e., the difference in length between any two divergence vectors) (Collyer & Adams 2007; Stuart *et al.* 2017). Small angles and small differences in length imply highly parallel evolution. With these summary statistics for (non)parallel evolution, we asked the following two questions:

1. What is the relative extent of phenotypic and genetic (non)parallelism at global versus regional scales?
2. What are the potential ecological and genetic drivers of phenotypic and genetic (non)parallelism at each scale?

## MATERIALS AND METHODS

### Field collections and environmental data

We used minnow traps to collect 39-40 lake and 40 stream stickleback, between May 21 and June 28, 2013, from each of six independent population pairs on Vancouver Island (the “regional” samples: Table S1A, Fig. S1A). We also used minnow traps or nets to collect 40 lake and 40 stream stickleback, between April 10 and August 8, 2014, from each of six other population pairs in North America and Europe (global samples: Table S1B, Fig. S1B). In all collections, we trapped the lake fish at least 100 m away from the junction with any stream (apart from Lake Constance, where the lake fish were captured at the mouth of the stream), and we trapped stream fish at least 100 m away from the junction with any lake. We did not retain any fish younger than one year or any obviously gravid females.

We euthanized the fish with an overdose of MS-222 fish anesthetic (Argent Chemical Laboratories, Redmond, WA), and removed the right pectoral fin, which was preserved in 95% ethanol for genetic analysis. We then fixed the fish in 10% neutral buffered formalin (VWR, Radnor, Pennsylvania). To facilitate gill raker counts, we stained the fish using alizarin red dye. To do so, we first soaked the fish in water for 24 hours, then stained the fish in a solution of alizarin red and 0.5% KOH for 24 hours, and then performed a second soak in water for 24 hours to remove excess stain. We then stored the fish in 40% isopropyl alcohol until they were processed further (see below), approximately four weeks later.

To characterize the environment at each site, we measured several habitat variables. Using the Google Earth Pro V7.3 software, we measured total lake area (m^2^), lake perimeter (m), and mean lake-stream elevation for each pair (m). Using measuring sticks and a tape meter, we also measured stream depth (cm), lake depth (cm), and stream width (cm) at each trap as well as stream flow (cm/s) using a flow meter.

### Morphological measurements

In the laboratory, we photographed the left and ventral surfaces of each fish against a grid marked at 1 cm intervals. The grid was placed at a constant distance from the lens of a Canon G11 digital camera, which was positioned in the same plane (using a surface level) as the grid. Before the photograph was taken, we removed the left pectoral fin of each fish and pinned it at maximum extension onto the same grid (for measurements of fin area, perimeter, and the length of fin rays). We also inserted small pins into the fish to help indicate otherwise cryptic homologous anatomical points for subsequent geometric morphometrics (Berner *et al.* 2009). After photographing, we measured several morphological traits (described below) with a focus on body shape and gill raker traits because repeated patterns of phenotypic divergence between lake and stream fish often involve these traits (Sharpe *et al.* 2008; Berner *et al.* 2008; Hendry *et al.* 2011; Berner *et al.* 2011; Kaeuffer *et al.* 2012; Oke *et al.* 2016; Moser *et al.* 2016). In addition, we dissected each fish to determine sex by gonad inspection.

From the photographs, we measured 25 linear trait distances (Table S2), as well as the area and perimeter of the pectoral fin using the image processing software *Fiji* (Schindelin *et al.* 2012). We standardized these 27 univariate measurements to a common standard body length with the allometric formula *M_S_ = M_0_ (L_S_ / L_0_)^b^*, where *M_S_* is the standardized trait measurement, *M_0_* is the unstandardized trait measurement, *L_S_* is the overall mean body size of all fish in a given analysis, and *L_0_* is the standard length of the individual (Elliott *et al.* 1994; Lleonart *et al.* 2000). Standard length is defined as the distance from the anterior tip of the upper jaw to the posterior end of the hyperal plate. The exponent *b* was calculated as the common within-group slope (Reist 1986) from a linear mixed-effect model regressing log_10_(*M_0_*) on log_10_(*L_0_*) with pair as the random factor.

We counted the number of gill rakers on the first right gill arch *in situ*, then removed the right gill arch and photographed it under a dissecting scope using an ocular micrometer and a Canon EOS Rebel T5 digital camera. To quantify gill raker length, we measured the curved distance along the centre of the three longest gill rakers. We then calculated raker spacing as 5/*l*, where *l* is the mm distance along the gill arch spanned by the five longest rakers. These four measurements (lengths of the three longest gill rakers, and gill raker spacing) were standardized to a common standard body length as described above.

To quantify overall geometric morphometric shape differences, we placed 19 homologous landmarks on the lateral photographs using the *TPSdig* software (Rohlf 2006) (Fig. S2). We then superimposed the 19 landmarks using a generalized Procrustes analysis (GPA) with *R* package *geomorph* (Adams & Otárola Castillo 2013). We performed this analysis for the global samples and the regional samples combined so that all fish would be positioned in the same shape space. These alignments resulted in 38 Procrustes residuals that described the shape differences between specimens unrelated to scale, rotation, or translation. The 38 principle components (PCs) derived from the 38 Procrustes residuals were then allometrically adjusted for centroid size and body depth using the above common within-slope approach (Reist 1986; Lleonart *et al.* 2000; Rolshausen *et al.* 2015). The first PC was found to describe shape differences due to vertical bending of the fish produced during storage. We therefore excluded this artefactual PC from further analyses.

### DNA extraction and restriction site associated DNA sequencing library construction

DNA was extracted from stickleback fin clips using the Wizard^®^ SV Genomic Purification kit (Promega^®^ Corp., Madison, USA) and quantified using Picogreen^®^ ds DNA assay (Thermo Fisher Scientific, Waltham, USA) on an Infinite^®^ 200 Nanoquant (Tecan Group Ltd. Männedorf, Switzerland). All samples were normalized to a dsDNA concentration of 15ng/µl, re-quantified, and pooled according to sampling location. Thus, we created 24 pools of 40 individuals each, consisting of 12 pools for each geographic scale (at each scale, six lake and six stream pools).

ddRAD PoolSeq libraries were prepared following a modified version of the Peterson *et al*. (2012) protocol. Briefly, pooled samples were first digested using the NLAIII and MluCI enzymes (New England Biolabs Inc., Frankfurt, Germany), individually barcoded using unique P1 flex-adapters (Peterson *et al.* 2012), and pooled together into a single library. Fragments of 371-416bp were extracted using a Pippen Prep^®^ 2% MarkerB 100-600bp cassette (Sage Science Inc., Beverly, USA). In post-size selection, we isolated fragments including at least one biotin-tagged P2 adapter, using streptavidin-couple beads (Dynabeads^®^ M-270, Thermo Fisher Scientific, Waltham, USA). The library was then PCR amplified in five separate aliquots (12 cycles, Phusion High Fidelity^®^ PCR kit, New England Biolabs Inc., Frankfurt, Germany). We used Agencourt^®^ AMPure XP beads (Beckman Coulter, Indianapolis, USA) to clean samples after each enzymatic reaction.

### Sequencing and ddRAD data processing

Barcoded ddRAD samples were sequenced on four lanes of an Illumina Hi-Seq 2500 for 100 bp paired-end (∼150M reads total), and demultiplexed using *STACKS* (Catchen *et al.* 2013) v. 1.13 (*process_radtags* -P, -p, -r, -i, --inline_index, --disable_rad_check). Resulting FASTA files were trimmed using *PoPoolation* (Kofler *et al.* 2011a) to eliminate low quality regions (--min-length 50 --quality-threshold 20). Reads from all 24 pools were aligned to the threespine stickleback genome (release-084, Ensembl.org) separately using *BWA mem* v. 0.7.13 (Li & Durbin 2009). We then used *SAMtools* (Li *et al.* 2009) v. 1.3.1 to convert the resulting sam files to sorted bam format, keeping only reads with mapping quality above 20 (*samtools view* -q 20). This led to an average of ∼53M reads per pool. A pileup file was then generated using *SAMtools* v. 1.3.1 (*samtools mpileup* -B), and filtered for completeness by keeping reads with a minimum coverage of five. We then converted the pileup file to a sync file using *PoPoolation2* (Kofler *et al.* 2011b) for further downstream analysis.

### Statistical analysis

All statistical analyses were performed in *R* 3.2.2 (R Core Team 2015).

#### Variation in environmental variables

To test whether variation in habitat characteristics was greater at the global than regional scale, we compared within-scale variation for each environmental parameter using Bartlett’s tests (Bartlett 1937). In addition, we calculated levels of among-lake and among-stream variation for each environmental variable at each scale separately. To calculate among-lake and among-stream variation at each scale, we computed ANOVAs at each scale separately with each environmental variable as dependent variable and watershed as predictor. Among-lake or among-stream variation was calculated by dividing the pair term sum of squares by the total number of pairs in each scale respectively.

#### Univariate and multivariate phenotypic analysis

To reduce the large number of univariate traits, we first performed a principal components analysis (PCA) on the size-standardized trait values for all samples (global and regional) combined. To consider phenotypic differences between lake and stream stickleback across pairs for the global and regional sample sets separately, we then used multivariate analysis of covariance (MANCOVA) models, with Wilks’ lambda (λ) as the test statistic. We tested for differences in specific trait sets for the regional and global samples separately using three different MANCOVA models, with dependent variables as: i) all 27 (size-standardized) univariate measurements, ii) the five gill raker traits only, and iii) the PCs derived from the 38 Procrustes residuals.

To understand the contribution to parallelism of a subset of individual traits likely important to lake-stream divergence (body depth, gill raker number, gill raker spacing, and mean length of the three longest gill rakers), we analyzed each of these traits separately using an analysis of covariance (ANCOVA) at each regional and global scale. The independent variables in every model were the fixed effects of sex, pair, and habitat (lake versus stream), along with all two-way interactions. Centroid size was also included as a covariate (along with its two-way interactions) to control for any remaining size effects.

#### Estimation of genome-wide population differentiation

We evaluated the genetic relationship within pairs by estimating the fixation index (*F*_ST_) for each lake-stream pairwise comparison using *PoPoolation2* (Kofler *et al.* 2011b), applying a number of stringent criteria (see below) to define genomic sites for analysis across the entire genome. The accuracy of allele frequency estimation of pooled individuals is highly dependent on sequence coverage (Schloetterer *et al.* 2014). Therefore, to reduce stochastic error and increase the accuracy of *F*_ST_ estimates, for each pairwise combination we used a minimum minor allele count of two across all pools, a high sequence coverage filter (between five and 500 within each pool), and a large 100-kb non-overlapping sliding window. This window size led to mean of 146.74 SNPs per window (± 90.29, median = 123). We visualized population structure by building neighbour-joining trees for both scales separately based on average pairwise *F*_ST_ values.

#### Evaluation of (non)parallelism at both geographic scales

We used two different approaches to estimate the extent of parallel evolution at the two geographic scales.

First, we compared the effect sizes of the habitat and habitat-by-pair interaction terms from the MANCOVA and ANCOVA analyses of phenotype (partial η^2^ and η^2^ values, respectively) (Oke *et al.* 2016; Stuart *et al.* 2017). A large effect size for the habitat term suggests consistent lake versus stream differences shared across pairs, i.e., parallel evolution. A large effect size for the pair-by-habitat interaction term, on the other hand, suggests idiosyncratic lake-versus-stream differences across pairs, i.e., non-parallel evolution. Thus, if the ratio of the habitat to the interaction effect size is greater in the pairs sampled at a regional scale as compared to the pairs sampled at a global scale, we infer that lake-stream divergence is more parallel at the regional scale.

Second, we calculated lake-to-stream divergence vectors for our phenotypic and genetic data. Each vector connects the multivariate lake mean to the multivariate stream mean and quantifies the magnitude and direction of lake-stream divergence for each population pair (Adams & Collyer 2009). Then, to describe the extent to which vectors diverged in parallel across pairs, we calculated: θ, the angle between any two vectors; L, the length of the vector for each pair; and ΔL, the difference in length between any two vectors. Strict parallel divergence between pairs would correspond to θ = 0 and ΔL = 0. Vector analysis for the phenotypic and genetic datasets follows Stuart *et al*. (2017) and is briefly described below:

i. Phenotypic: For each pair, for each trait (Table S2), we ran a t-test comparing the distribution of a trait in a lake to that in its adjoining stream. Concatenating each t-statistic resulted in a 1xN vector of scale-standardized lake-stream differences, thereby constituting our lake-stream divergence vector. We then calculated angles between phenotypic divergence vectors (θ_P_) as the arc-cosine of the Pearson correlation for each vector pair. L_P_ was the multivariate Euclidian length of each vector, and ΔL_P_ was the difference in length between any two divergence vectors. We tested for differences in θ_P_ and ΔL_P_ between global and regional scales using a two-tailed t-test.
ii. Genetic: Using the sync file of allele frequencies extracted with *PoPoolation*, we first calculated the frequency of the minor allele at each SNP. We then used PCA (*R prcomp*) to reduce our large SNP dataset into a smaller number of quantitative axes of genetic differentiation. We saved populations’ scores of all the resulting 24 axes. We calculated the centroid for each lake and each stream, for these 24 axes, to obtain vectors in genotypic space. Thus, each row of the resulting matrix represents the estimate of genetic divergence between the two habitats of the same pair, thereby constituting the genetic lake-stream divergence vector for each pair. First, we performed this analysis on a set of ‘outlier’ SNPs whose lake-stream *F*_ST_ values fell within the top 5% of estimated values for each of the 12 lake-stream pairs, thus representing markers putatively linked to selected loci (175,302 outlier loci). To test for differences in genetic divergence between scales, we performed the genetic vector analysis as described above and across both scales to obtain measures of θ_OUTLIERS_ and ΔL_OUTLIERS_. We tested for differences in θ_OUTLIERS_ and ΔL_OUTLIERS_ between global and regional scales using a two-sided t-test. Second, we calculated θ, L, ΔL by excluding these outlier SNPs to obtain a set of markers likely to represent neutral population genetic processes (327,291 neutral loci - hereafter θ_G_, L_G_, and ΔL_G_). We tested for differences in θ_G_ and ΔL_G_ between global and regional scales using a two-sided t-test. Values of θ_G_, L_G_, and ΔL_G_ were used further to investigate potential constraints on divergence (see below). Finally, we considered whether neutral processes could be responsible for phenotypic divergence by testing for a linear relationship between θ_P_ values and averaged between-watershed neutral *F*_ST_ at each scale separately using linear models with *F*_ST_ as predictor and θ_P_ as response variable.

Third, we verified whether watersheds share the same genetic architecture underlying adaptive traits by calculating the proportion of shared outlier loci across population pairs. At both geographic scales, we counted the total number of outlier loci and calculated the proportion of shared outlier loci over one or multiple watersheds.

Finally, because it is possible that some environmental variables could be confounded with geographic scale (e.g., the lakes and streams that we sampled at the global scale were on average larger than those sampled at the regional scale), we tested the effect sizes of our measured environmental variables relative to geographic scale in models where both forms of information acted as predictors for phenotypic or genetic divergence.

## RESULTS

### Variation in environmental variables

Overall, environmental variation is higher at the global scale. For all environmental variables (in both lake and stream habitats), variation among watersheds was significantly greater at the global than regional scale (Table 1, Fig. S3). Similarly, among-lake and among-stream variation was greater at the global than at the regional scale (Fig. S4).

**Table 1.** Within-scale mean and standard deviation in environmental variables. Results from Bartlett’s tests of homogeneity of variances are presented with K-squared (*K^2^*), degrees of freedom (df), and *P*-values (*P*-value). *P*-values < 0.05 are in bold.

|  | Mean |  | Standard deviation |  | Bartlett test |  |  |
| --- | --- | --- | --- | --- | --- | --- | --- |
| | Regional | Global | Regional | Global | $K^2$ | df | $P$-value |
| Lake area (m<sup>2</sup>) | 8.50<sup>e+05</sup> | 1.13<sup>e+08</sup> | 7.16<sup>e+05</sup> | 2.25<sup>e+08</sup> | 45.98 | 1 | **<0.001** |
| Lake perimeter (m) | 6.19<sup>e+03</sup> | 7.19<sup>e+04</sup> | 5.84<sup>e+03</sup> | 1.00<sup>e+05</sup> | 19.56 | 1 | **<0.001** |
| Lake-stream elevation (m) | 106.33 | 100.17 | 72.80 | 159.75 | 2.56 | 1 | 0.109 |
| Lake depth (cm) | 14.02 | 33.25 | 12.93 | 35.96 | 3.84 | 1 | **0.004** |
| Stream depth (cm) | 73.14 | 75.37 | 738.62 | 4438.49 | 3.25 | 1 | 0.070 |
| Stream width (cm) | 505.62 | 2016.57 | 122.96 | 2883.69 | 22.39 | 1 | **<0.001** |
| Stream flow (cm/s) | 3.72 | 131.60 | 5.16 | 130.22 | 23.06 | 1 | **<0.001** |

### Univariate and multivariate phenotypic analysis

Consistent with previous work, we observed variable levels of (non)parallelism in univariate and multivariate traits. The first and second PCs for the 27 univariate measurements explained 24.7% and 17.5% of the total variance, respectively, and every trait except the length of the dorsal fin loaded negatively on the first axis. The within-pair difference between lake and stream fish on PC1 was always in the same direction, indicating that any habitat-by-pair interaction on PC1 was due to differences in the magnitude of divergence, rather than the direction of divergence (Fig. 1). The MANCOVA models revealed significant effects of habitat, pair, and their interaction on all sets of traits at both regional and global scales (Table S3).

**Figure 1:**
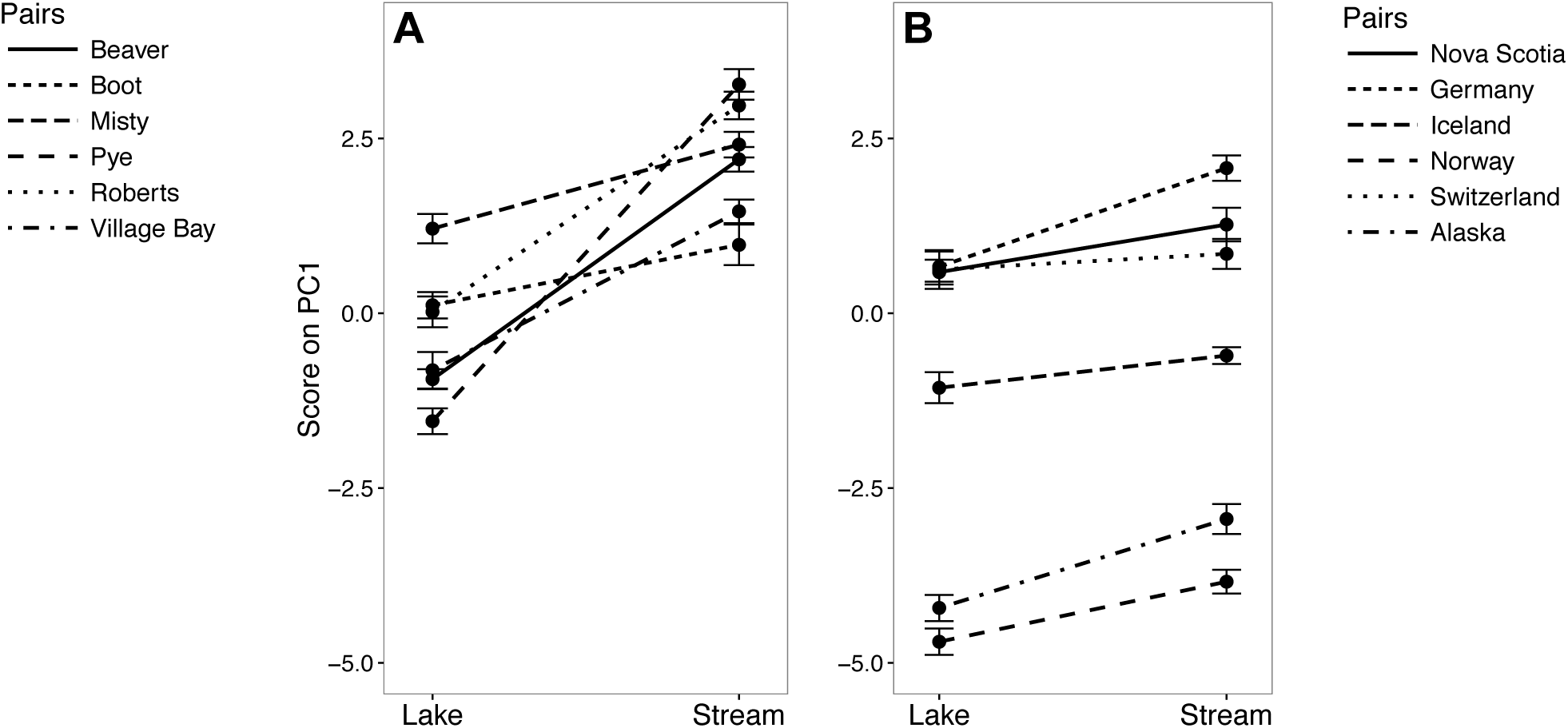
Mean and standard error of score on PC1 of the 27 univariate measurements for regional (A) and global (B) lake-stream pairs. The direction of phenotypic divergence is consistent but the magnitude of phenotypic divergence varies.

ANCOVAs for individual traits revealed that the effects of habitat, pair, and their interaction were always significant at both the regional and global scales (Table S4). Specifically, stream fish were generally deeper bodied than lake fish for the regional pairs, often with similar magnitudes of difference. However, in the global pairs, stream fish were sometimes not deeper bodied than their lake counterparts (Fig. 2A, B). Lake fish also generally had more and longer gill rakers than stream fish (Fig. 2C-F), although this difference was more pronounced at the regional scale. Gill raker spacing was greater in stream fish at the regional scale but not at the global scale (Fig. 2G, H).

**Figure 2:**
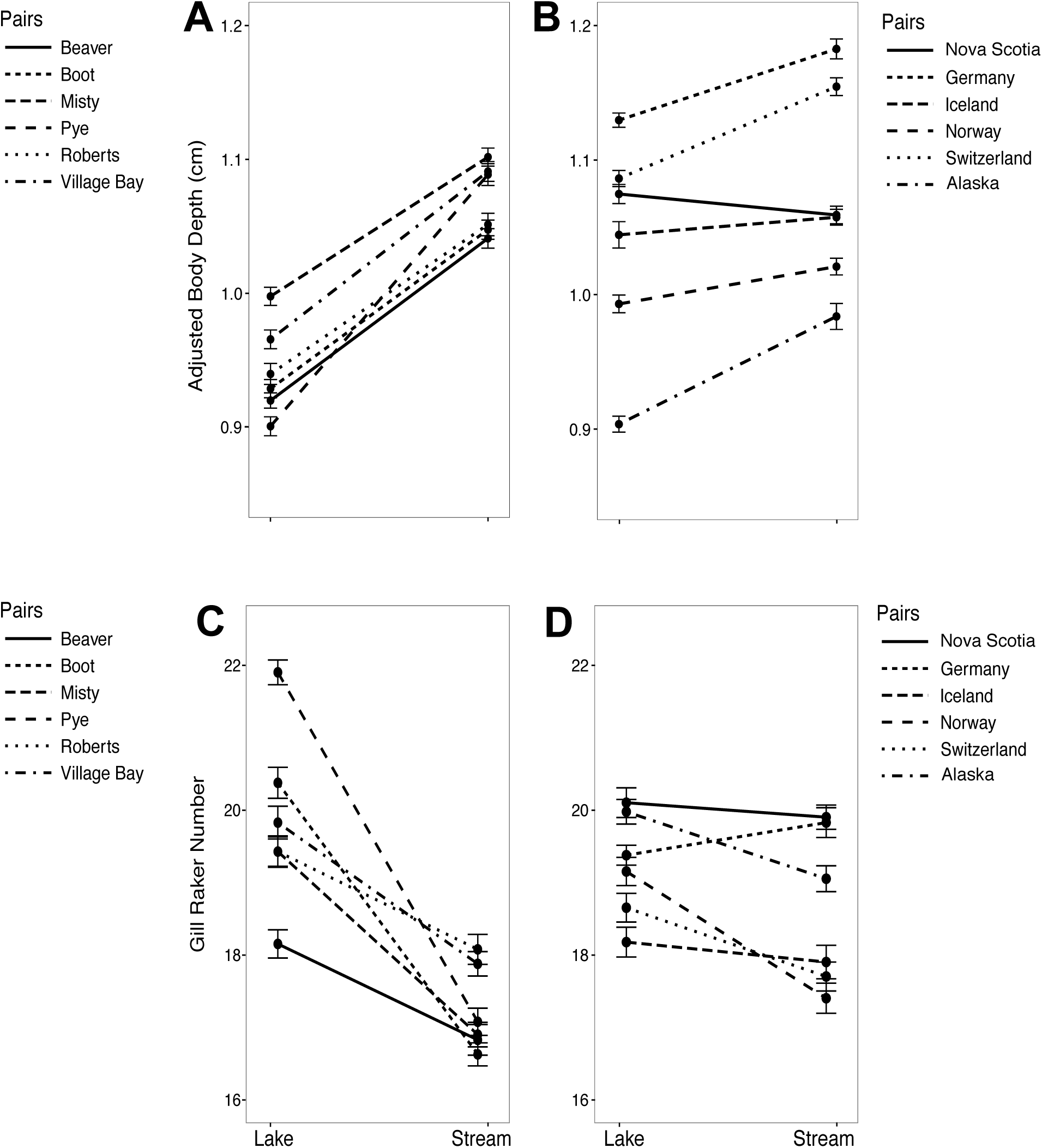
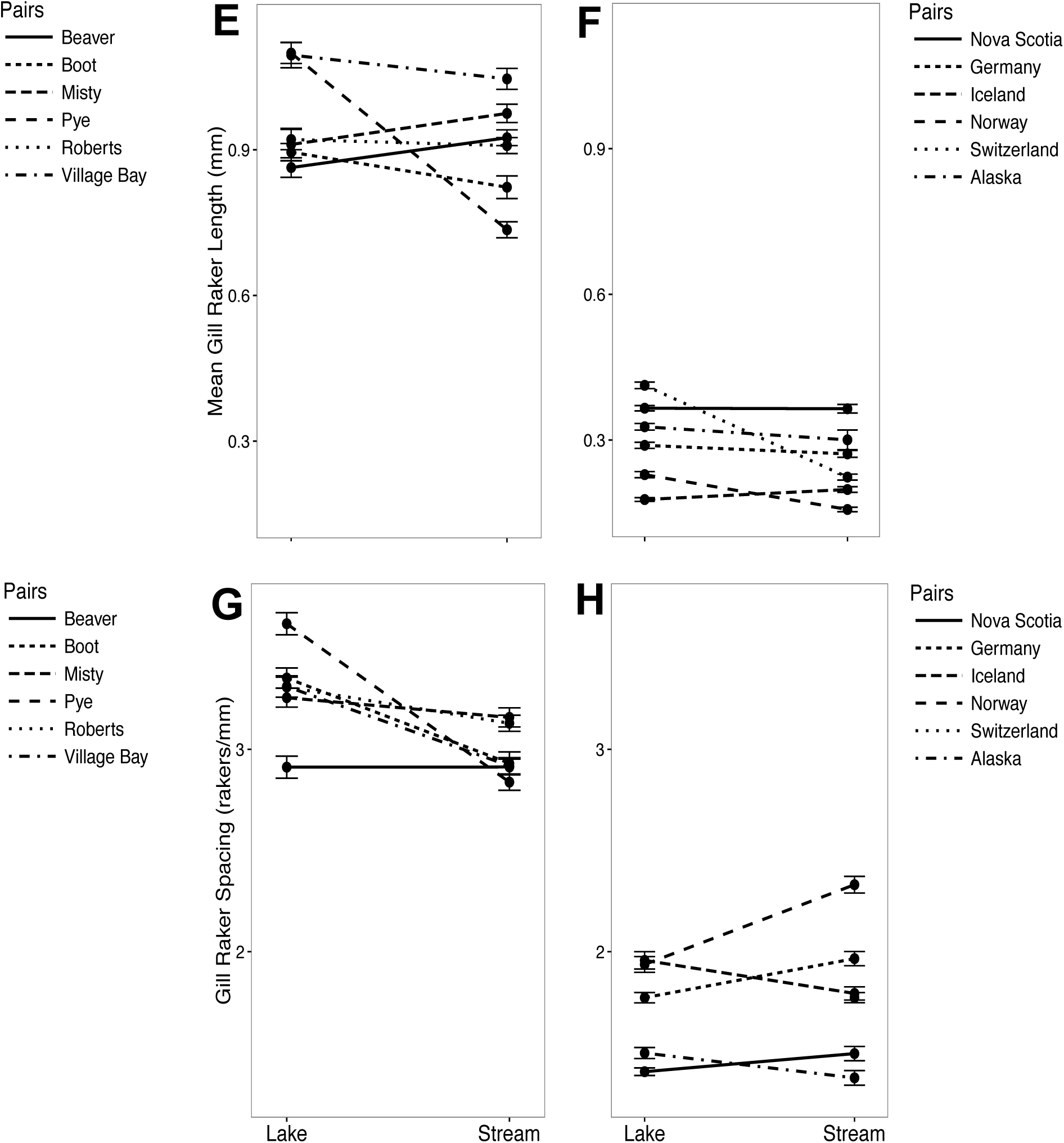
Mean and standard error of body depth (A, B), number of gill rakers (C, D), mean gill raker length (E, F) and gill raker spacing (G, H) for regional (left panels) and global (right panels) lake-stream pairs.

### Evaluation of (non)parallelism at both geographic scales

#### Differences in effect sizes of multivariate models

Significant differences in effect sizes from the MANCOVAs and ANCOVAs revealed that phenotypic parallelism was greater at the regional than at the global scale. Indeed, the ratio of the habitat term to the habitat by pair interaction term was always greater at the regional than at the global scale – across all models. In the MANCOVA models, the averaged ratio was 3.2 at the regional scale and 1.5 at the global scale. In the ANCOVA models, the averaged ratio was 6.7 at the regional scale and 0.3 at the global scale (Table 2, Fig. 3; see Tables S3 and S4 for more detailed results about these models). This finding suggests that lake-stream divergence in phenotypic traits was more parallel at the regional than global scale.

**Figure 3:**
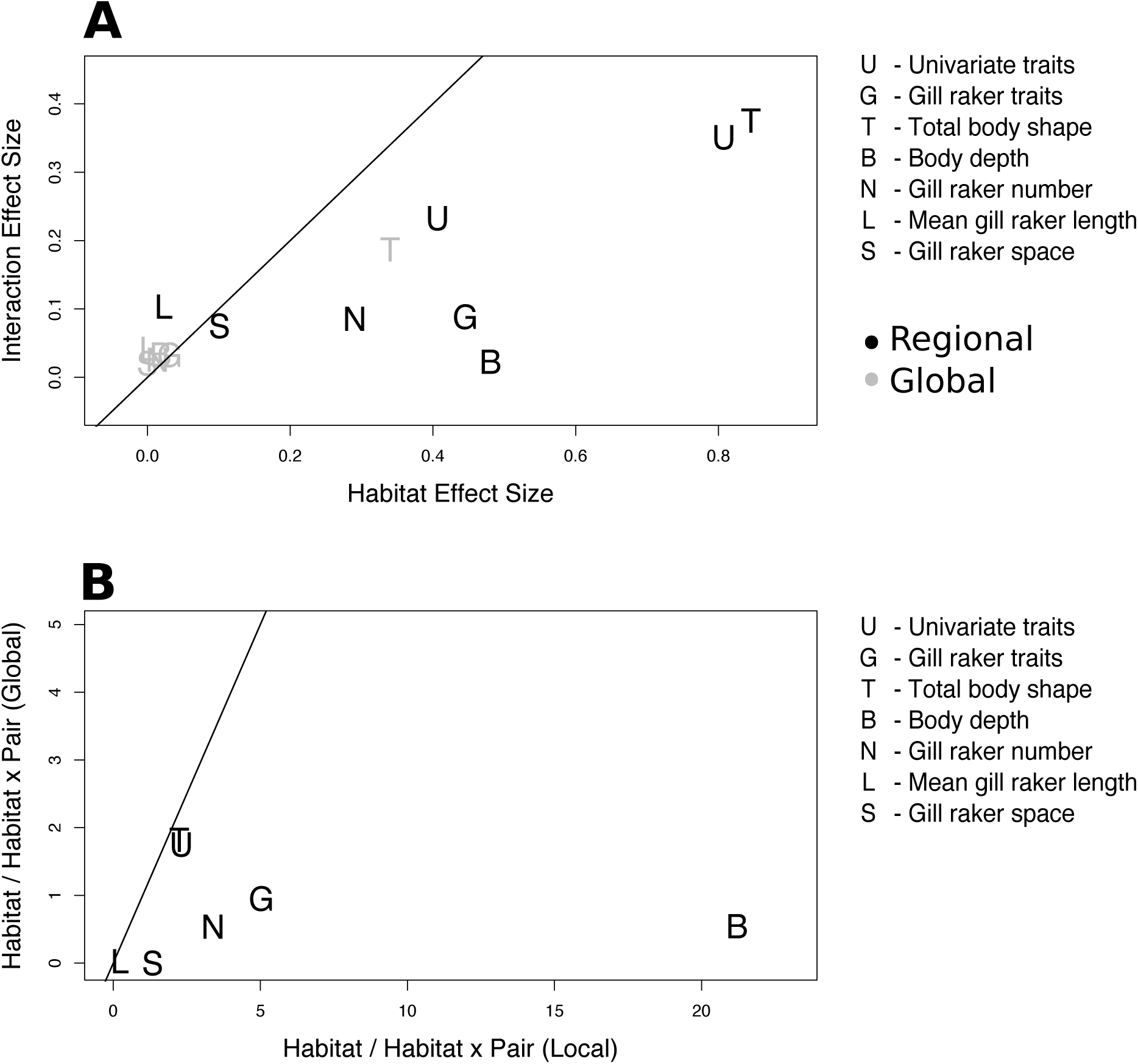
Phenotypic parallel divergence is higher in regional versus global comparisons. (A) Effect size of the habitat-by-pair interaction term versus effect size of the habitat term for regional and global pairs. (B) Ratio of the habitat effect size to the habitat-by-pair interaction effect size for regional pairs (x-axis) versus global pairs (y-axis) for all traits examined. In (A) and (B) effect size measured as η^2^ for single traits and partial η^2^ for multivariate traits. In (A), traits found below the 1:1 line show more parallelism (large, consistent effect of habitat) than non-parallelism (large, but inconsistent effect of habitat). Traits above the line show the opposite pattern. In (B) traits found below the 1:1 line show greater parallelism in regional than in the global pairs.

**Table 2.**
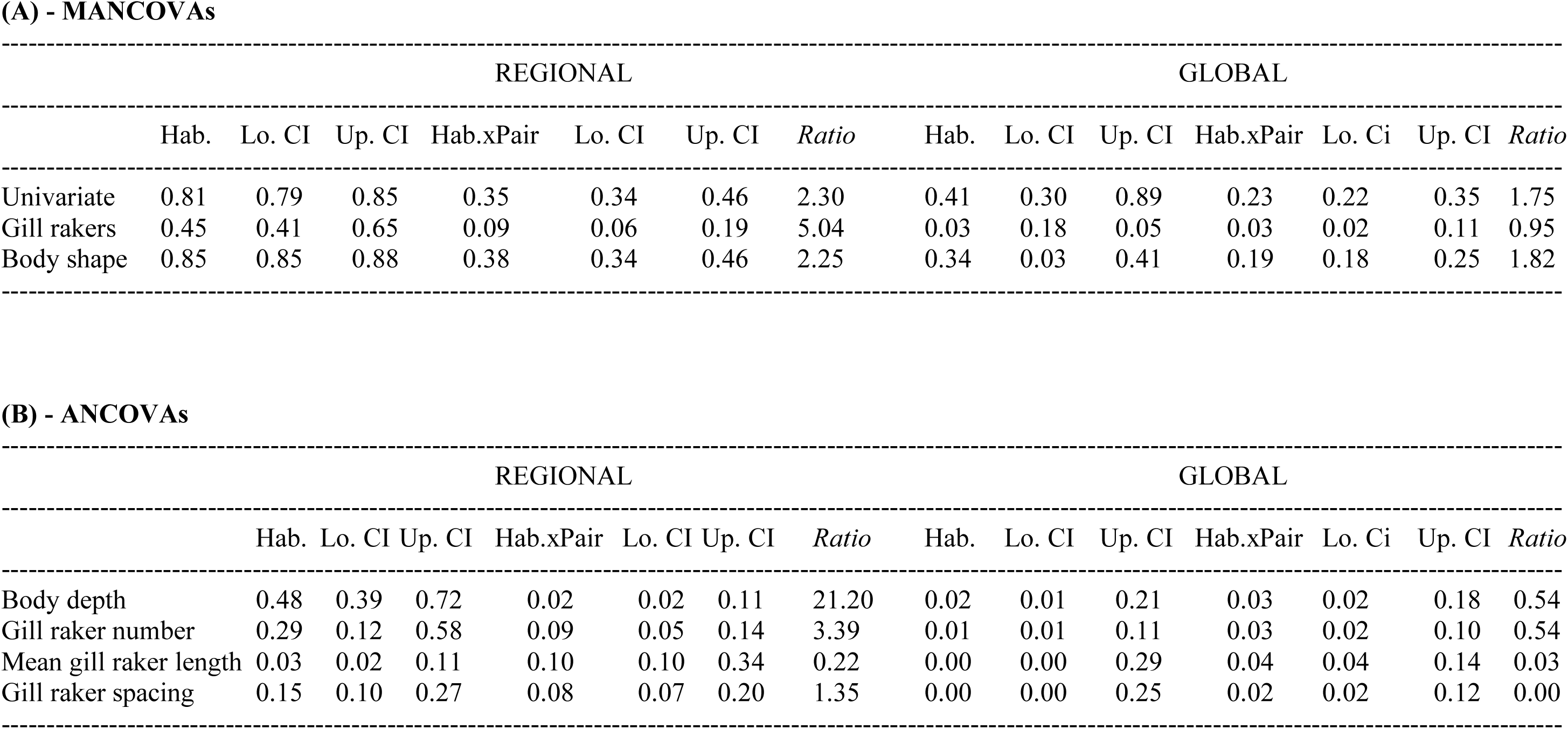
Results from MANCOVAs on the multivariate traits (A) and ANCOVAs on single traits (B) at both regional and global scales. Effect sizes (Partial η^2^) and their lower and upper 95% confidence limits (Lo.CI; Up.CI) of the habitat (Hab.) and habitat by pair interaction (Hab. x Pair) as well as their ratios (Ratio: Hab./Hab. x Pair) are reported.

#### Vector analysis

*F*_ST_ extracted from the sliding window analysis revealed that lake-stream populations in each pair were each other’s sister taxon, regardless of the geographic scale of comparison (Table S5, Fig. S5). This pattern is consistent with evolutionarily independent lake-stream divergence in each pair, although ongoing gene flow could also contribute to this pattern.

Overall, we found phenotypic divergence to be more parallel at the regional than global scale. Multivariate vector analysis of the phenotypic data revealed a continuum of lake-stream parallelism at both scales (Tables S6 and S7). The mean angle between any two phenotypic vectors (θ_P_) was 61.63° ± 12.24sd at the regional scale and 90.17° ± 15.98sd at the global scale (Fig. 4A; *t* = 5.49, df = 26.23, *P* = 8.97e^-06^). In addition, phenotypic vector lengths, which measure the magnitude (but not direction) of lake-stream divergence, also exhibit greater similarity among regional pairs than among global pairs. Pairwise comparisons of vector lengths (ΔL_P_) averaged −2.85 ± 10.57sd locally, and 6.52 ± 7.98sd globally (Fig. 4C; t = 3.16, df = 14.00, *P* = 0.007).

**Figure 4:**
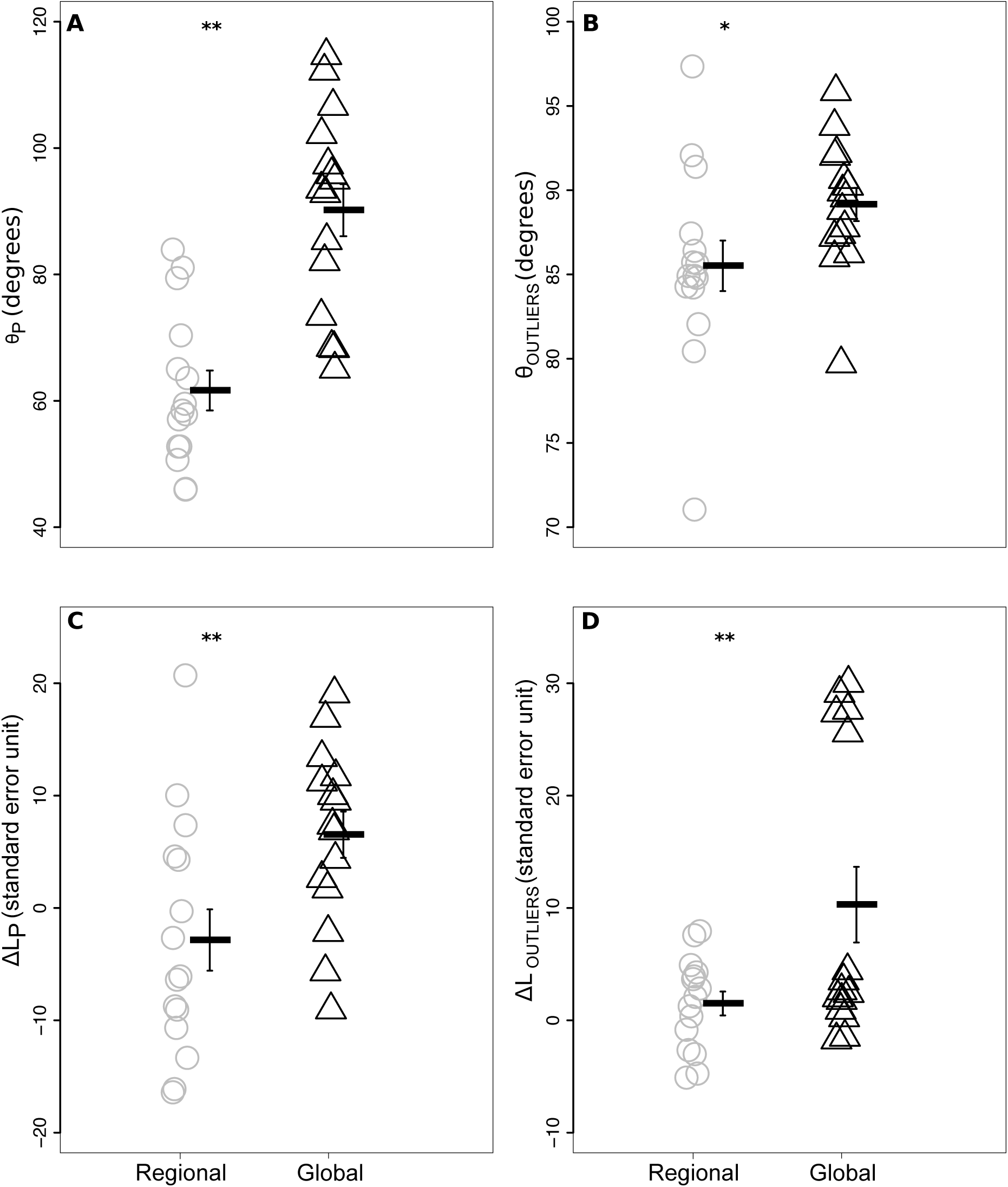
Pairwise angles between phenotypic vectors (Panel A, θ_P_), outlier genetic vectors (Panel B, θ_OUTLIERS_), as well as differences in vector length between phenotypic vectors (Panel C, ΔL_P_) and outlier genetic vectors (Panel D, ΔL_OUTLIERS_) at regional (grey circles) and global (black triangles) scales. Overall mean values are represented by a straight line (±SE). Significance in mean differences is represented by two stars (*P* < 0.01) or one star (*P* < 0.05).

At the genetic level, divergence was significantly more parallel at the regional than global scale, but only for outlier loci: The mean angle between any two outlier genetic vectors (θ_OUTLIERS_) was 85.51° ± 5.81sd at the regional scale and at 89.15° ± 3.83sd the global scale (Fig. 4B; t = 2.02, df = 24.21, *P* = 0.04). The mean angle between any two neutral genetic vectors (θ_G_) was 89.32° ± 2.72sd at the regional scale and 90.03° ± 1.81sd at the global scale (t = 0.85, df = 24.36, *P* = 0.41). Genetic vector lengths were significantly smaller at the regional than global scale at both outlier and neutral loci. However, at outlier loci, genetic vector length differences were primarily driven by one population (Norway, Table S9D). Outlier vector lengths (ΔL_OUTLIERS_) differed by a mean of 1.49 ± 4.12sd and 10.29 ± 13.05sd at the regional and global scales, respectively (Fig. 4D; t = 2.49, df = 16.75, *P* = 0.02). Neutral genetic vector lengths (ΔL_G_) differed by a mean of 0.46 ± 1.54sd and 6.12 ± 6.77sd at the regional and global scales, respectively (t = 3.16, df = 15.44, *P* = 0.006). Finally, θ_P_ and averaged between-watershed neutral *F*_ST_ were not significantly associated at either scale (regional: t = −0.88, df = 13, *P* = 0.39, *R^2^* = −0.24; global: t = 1.12, df = 13, *P* = 0.28, *R^2^* = 0.29).

If different watersheds share the same genetic architecture underlying adaptive traits, we would expect outlier loci under selection to overlap across population pairs when lake-stream divergence proceeds in parallel. However, 63% of the outlier loci were unique to a specific watershed across all watersheds, whereas 61% and 77% of outliers were unique to a specific watershed at the regional and global scales, respectively (21% and 4% were shared across a minimum of two watersheds at the regional scale and global scale, respectively). As an overall summary, comparisons of vector pairs showed less parallelism at the global scale than at the regional scale, at both the phenotypic and genetic outlier levels; for neutral genetic loci, (non)parallelism was similar at both scales. Importantly, perfect parallel divergence is non-existent at both phenotypic and genetic levels and at both geographic scales.

Finally, and with only a few exceptions (lake depth and stream depth with ΔL_P_, stream flow with θ_P_), environmental variables were not associated with divergence metrics (Table S10). Moreover, in only one case (lake depth and ΔL_P_) was the effect size of an environmental variable greater than that of geographic scale. Thus, the effect of geographic scale on divergence between lake and stream pairs appears to be mostly independent of any consistent environmental differences between the scales.

## DISCUSSION

We assessed the influence of geographic scale on the repeatability of adaptive radiation by quantifying levels of phenotypic and genetic parallelism in lake-stream population pairs of threespine stickleback. In agreement with previous work conducted over restricted spatial scales (Lavin & McPhail 1993; Hendry & Taylor 2004; Berner *et al.* 2008; 2009; Kaeuffer *et al.* 2012; Stuart *et al.* 2017), we found a continuum of phenotypic and genetic parallelism, with lake-stream divergence being more pronounced for some traits, and in some population pairs, than in others. At the phenotypic level, parallelism was especially notable in body depth and gill raker number – also consistent with previous work. When comparing between geographic scales, parallelism in phenotypes and genetic outliers was greater at the regional scale (Vancouver Island) than at the global scale (Europe and both sides of North America). In contrast, parallelism was similar across scales for neutral genetic loci. We suggest that the observed disparities in phenotypic and genetic parallelism between scales are likely the result of differences in environmental variation, selection pressures, and genetic architecture of traits between scales. We develop these ideas further in the text below.

### Phenotypic (non)parallelism

Univariate, multivariate, and vector analyses all revealed that phenotypic parallelism was greater at the regional than the global scale (Fig. 4A, C). This difference between scales was especially pronounced in specific morphological traits, such as body depth and gill raker length, that are known to have a strong heritable basis and that have been previously implicated in adaptive parallel evolution (Sharpe *et al.* 2008; Berner *et al.* 2008; Hendry *et al.* 2011; Berner *et al.* 2011; Kaeuffer *et al.* 2012; Miller *et al.* 2014; Oke *et al.* 2016; Moser *et al.* 2016).

Lake-stream body depth divergence was highly parallel in that stream fish always had deeper bodies than lake fish; yet the level of this parallelism was almost always greater at the regional than the global scale (Figs. 1-4). A similar pattern emerged for gill raker number, with strong parallelism evident in lake-stream divergence at the regional scale, consistent with previous studies (Berner et al. 2008; 2009; Kaeuffer et al. 2012; Stuart et al. 2017). Although lake-stream parallelism in these traits was also present at the global scale, it was less pronounced than at the regional scale (Table 2, Fig. 3). For instance, also in accordance with previous studies (Berner et al. 2008; Kaeuffer et al. 2012), gill raker spacing and length exhibited low levels of parallelism (Fig. 2E-H), yet the parallelism that was present in these traits was greater at the regional than at the global scale (Fig. 3B).

With environmental variation across watersheds being greater at global than regional scales (Figs. S3, S4), our data are consistent with the hypothesis that reduced environmental variation among watersheds at smaller spatial scales enhances (or enables) stronger parallel evolution. For instance, divergent selection related to diet, foraging mode, and swimming performance has been implicated in variation in body shape and gill rakers between lake and stream stickleback. In lakes, sticklebacks generally feed on small limnetic prey (Berner *et al.* 2008; 2009; Kaeuffer *et al.* 2012), leading to selection for an elongated body shape (Webb 1984; Walker 1997) and longer/more numerous gill rakers (Bentzen & McPhail 1984). In streams, stickleback commonly feed on benthic prey, leading to selection for deeper bodies and shorter/less numerous gill rakers (Bentzen & McPhail 1984; Berner *et al.* 2008; 2009; Hendry *et al.* 2011; Kaeuffer *et al.* 2012). Body depth and gill raker number are both known to be strongly heritable (Hermida *et al.* 2002; Aguirre *et al.* 2004; Oke *et al.* 2016; McPhail), and as such, the stronger parallelism of these traits at regional than at global scales presumably reflects greater determinism of selection owing to more consistent ecological differences between these habitat types across replicate watersheds on smaller spatial scales.

### Genetic (non)parallelism

Like phenotypic parallelism, genetic parallelism at outlier loci was more pronounced at the regional than the global scale, especially in angles (θ_OUTLIERS_). In contrast, for neutral loci, genetic parallelism was similar at the two geographic scales. Why might we observe greater parallelism at the regional versus global scales for outlier genotypes? One explanation is that these outliers are associated with the underlying genetic basis of adaptive phenotypes, which – as described above – show greater parallelism at the regional than the global scale. However, phenotypic parallelism does not necessarily lead to genetic parallelism if different genetic changes are capable of attaining similar phenotypic outcomes (Kautt *et al.* 2012; Conte *et al.* 2012; Westram *et al.* 2014; Le Moan *et al.* 2016). The greater parallelism of outlier loci at regional than global scales might thus reflect more similar genetic architectures and higher levels of shared standing genetic variation available to selection at smaller geographical scales (Barrett & Schluter, 2008; Agrawal & Stinchcombe, 2009; Flint & Mackay, 2009; Terekhanova *et al.*, 2014; Saltz *et al.*, 2017; Llaurens *et al.*, 2017; Bassham *et al.*, 2018). In support of this idea, we found that Vancouver Island lake-stream pairs shared more outlier loci than did lake-stream pairs at the global scale (21% of outliers were shared across a minimum of two pairs at the regional scale but only 4% of outliers were shared across a minimum of two pairs at the global scale). Note however that the interpretation of the above results might be more specific to the direction of divergence rather than the magnitude. Indeed, outlier genetic vector length differences were mostly driven by one population (Norway, Table S9D). Thus, rather than the magnitude of genetic divergence, differences between scale might mostly be driven by the relative weight of specific outlier loci in contributing to lake-stream divergence.

In contrast to the outlier loci, we detected no differences between geographic scales in the levels of (non)parallelism at neutral loci – at least when quantified using the *direction* (θ_G_) of genetic vectors. However, neutral genetic vectors were somewhat more similar in *magnitude* (ΔL_G_) at the regional than the global scale. This last result might suggest a role for gene flow in reducing divergence between watersheds over smaller spatial scales – that is, they are less genetically “independent” not only in their colonization but also in their contemporary genetic connections (Stuart *et al.* 2017; Berner & Roesti 2017; Haenel *et al.* 2018). It is also possible that these patterns could be driven by a difference in the effect of drift in shaping the extent of lake-stream neutral genetic divergence between scales. Nonetheless, the lack of differences in θ_G_ between scales, and the lack of a relationship between θ_P_ and the average between-watershed neutral *F*_ST_ at either scale, imply that neutral processes are unlikely to be responsible for phenotypic divergence at either scale.

Our conclusion that parallelism is stronger at the regional than the global scale is based on only a single region (Vancouver Island). Thus, the alternative conclusion might be that Vancouver Island lake-stream stickleback are simply more parallel than lake-stream stickleback in other locations. Unfortunately, few of the regions we examined contain numerous lake-stream pairs, thus precluding a formal comparison. However, several studies have compared multiple lake-stream pairs within some regions – frequently revealing strong parallelism. For instance, strong within-region lake-stream parallelism has been argued for Ireland (Ravinet *et al.* 2013), Iceland (Lucek *et al.* 2014b), Switzerland (Lucek *et al.* 2014b), and Haida Gwaii (Deagle *et al.*, 2012). Thus, while it is possible that parallelism is stronger on Vancouver Island than in other regions (Berner *et al.*, 2010), it is clear that global lake-stream parallelism is even lower – as revealed here.

### Constraints on parallel evolution at regional versus global scales

Lake-stream pairs at both scales showed relatively low levels of parallel evolution, especially for multivariate traits. Indeed, the mean angles between any two phenotypic vectors (θ_P_) were 61.63° ± 12.24 sd and 90.17° ± 15.98 sd at the regional and global scales, respectively. Similarly, the mean angles between any two genetic outlier vectors (θ_OUTLIERS_) were 85.51° ± 5.81 sd at the regional scale and 89.15° ± 3.83 sd at the global scale. Multiple factors can restrict the level of parallel evolution at both scales (Bolnick *et al.* 2018). For instance, a number of ecological variables that differ between lakes and streams are known to affect the degree of stickleback divergence, including diet (Berner *et al.* 2008; Kaeuffer *et al.* 2012; Ravinet *et al.* 2013), parasite load (Eizaguirre *et al.* 2011), flow regimes (generally higher in streams: Jiang *et al.* 2015), and predators (predatory fish and birds likely more common in lakes: Reimchen *et al.* 1985; Reimchen 1994). However, the direction and magnitude of lake-stream divergence in these factors is not always consistent (Berner *et al.* 2008; Kaeuffer *et al.* 2012; Stuart *et al.* 2017). Parasite communities are not universally more diverse in lakes than streams (Feulner *et al.* 2015), flow can be low in many stream reaches (Moore *et al.* 2007), and fish and bird predators vary dramatically among different lakes – and among different streams (Moodie & Reimchen 1976). Thus, variation across watersheds in the consistency of the ecological differences between lake and stream habitats could disrupt patterns of parallel evolution. We found this type of among-watershed environmental variation to be highest at the global geographic scale, and suggest that this non-parallel ecological divergence is likely to be a strong contributing factor to the non-parallel phenotypic and genetic divergence. However, it is noteworthy that univariate traits, such as body depth and gill raker number, were highly parallel in comparison to multivariate traits. One possible explanation is that “simple” univariate traits generally experience more parallel selection than “complex” multivariate traits for which multiple evolutionary solutions are possible to the same ecological challenge (Thompson *et al.* 2017).

At the genetic level, several factors can constrain the level of parallel evolution. Parallel evolution at loci associated with ecologically-relevant phenotypes is likely shaped by environmental heterogeneity for the same reasons as those mentioned for phenotypes above (Orr 2005). Physically proximate populations are also be expected to be more closely related genetically (both because of colonization and gene flow), and therefore could be expected to display greater levels of parallelism due to sharing a more similar molecular basis of phenotypes (Barrett & Schluter 2008; Conte *et al.* 2012). For instance, with partially-shared pools of standing genetic variation, closely related populations might have more similar genetic solutions to similar adaptive problems (Barrett & Schluter 2008). The Vancouver Island populations in our study likely diverged from a marine ancestor at a similar time ∼ 5,000 years ago (Stuart *et al.* 2017). In contrast, the global scale populations are estimated to have diverged at quite different times: as early as 5,000-15,000 years ago in Alaska (Reger & Pinney, 1996; Cresko *et al.*, 2004) to hundreds to thousands of years in Europe and Iceland (Moser *et al.* 2012; Lucek *et al.* 2014b; Roesti *et al.* 2015). This discrepancy between scales in divergence times might also explain the observed differences in parallel evolution; for the reasons outlined above, populations with similar divergence times might be more parallel than populations with diverse divergence times. Gene flow is another factor known to effect adaptive divergence (Slatkin, 1987; Lenormand, 2002; Garant *et al.*, 2007) and therefore also evolutionary (non)parallelism. On the one hand, populations exchanging genes will become more genetically similar, which can promote parallelism. On the other hand, gene flow between populations in distinct environments can impede regional adaptation (Garant *et al.*, 2007). Thus, gene flow within watersheds containing populations inhabiting distinct lake and stream habitat types should generally constrain their adaptive differentiation – as has frequently been inferred for lake-stream stickleback (Hendry *et al.*, 2002; Moore *et al.*, 2007; Berner *et al.*, 2009; Stuart *et al.*, 2017). If levels of gene flow are high within some watersheds and low in others, parallelism could be restricted, in particular as reflected by ΔL (Stuart *et al.* 2017). We detected lower ΔL_P_ and ΔL_G_ at regional than global scales, suggesting that the role of within-watershed gene flow in constraining adaptive differentiation could be more consistent at smaller spatial scales. However, ΔL_OUTLIERS_ was not different between the regional and global scales, which could imply similar levels of within-watershed gene flow across scales for outlier loci. The differences in ΔL_P_ may then reflect either differing levels of phenotypic plasticity between scales (Oke *et al.* 2016), or that our genetic markers have not included the loci associated with the traits experiencing parallel adaptive divergence between habitats, which might show distinct patterns of admixture to those described here (Nosil *et al.* 2009).

### Implications for adaptive radiation

Threespine stickleback are a remarkable example of adaptive radiation: they show dramatic phenotypic and evolutionary divergence from very small to very large geographical scales (Bell & Foster, 1994; McKinnon and Rundle 2002). Moreover, most of this divergence has played out over just a few thousand generations; and, in some cases, impressive freshwater divergence from a marine ancestor has evolved in less than fifty years (Kimmel *et al.*, 2012; Lescak *et al.*, 2015; Bassham *et al.*, 2018). At the same time, different stickleback populations in the adaptive radiation can still interbreed in nearly all cases (Hendry, 2009); and they show at most two forms in a single location (but see: Hippel von & Weigner, 2004). Hence, the applicability of the stickleback radiation to inferences about other classic adaptive radiations should be limited to the early stages of those radiations. Yet this potential limitation is also a strength in some respects; by capturing multiple replicates of the very early stages of adaptive radiation, we can more easily draw inferences about the processes that promote and constrain adaptive radiation – because we are removed from concern over confounding effects of non-causal factors that accumulate after the radiation is largely complete.

From this perspective, we found that, when replicate adaptive radiations are farther apart in space, and therefore are more likely to be independent, they show greater dissimilarities as reflected in lower phenotypic and genetic parallelism. We suggest that this scale-dependent repeatability is primarily driven by ecological rather than genetic factors. That is, the set of divergent environments in a given location will differ more from the same set of divergent environments in another location when those locations are situated farther apart in space. Indeed, the largest differences between scales appears to derive from phenotypes and genetic outliers, as opposed to neutral genetic markers. Further effort should be directed toward exploring both the ecological and genetic causes of scale-dependent determinism.

## ACKNOWLEDGEMENTS

This work would not have been possible without help from Katie Peichel, Elena Motivans, Mingsha Zhou, Susannah Halbrook, Matt Dubin, and Dario Moser in sampling sticklebacks. Thanks also to Mehvish Bukhari and Louis Astorg for their assistance with gill raker measurements. Thank you to Daniel Berner for providing samples from the Swiss pairs as well as environmental data and for providing a friendly review of the manuscript. Thanks to the Mullaney family for their assistance with catching Nova Scotia sticklebacks. A SEED grant from the Quebec Centre for Biodiversity Science to DH, Natural Sciences and Engineering Council (NSERC) of Canada Discovery Grants to RDHB and APH, and a Canada Research Chair Grant to RDHB provided financial assistance. The research was supported by NSF grants DEB-1144773 (DIB) and DEB 1456462 (YES & DIB). The Vancouver Island data were collected with funds from NSF grant DEB-1144773 to DIB.

## DATA AVAILABILITY

We have deposited the primary data underlying these analyses as follows:

- Raw phenotypic data and sync file of allele frequencies: Dryad

## AUTHOR CONTRIBUTIONS

AP, DH, APH, and RDHB designed the study. YS and DIB provided sequence and environmental data from the regional pairs. FAvH, MK, TK, SS and BKK provided fish samples and collected environmental data from the global pairs. AP and DH extracted sequence data from the global pairs and phenotypic data from both scales. AP and DH analyzed the data. AP and DH wrote the manuscript with input from all coauthors.

