## Supplementary data for "Repeatability of adaptive radiation depends on spatial scale: regional versus global replicates of stickleback in lake versus stream habitats"

### SUPPLEMENTARY MATERIAL

**Table S1**. Sampling sites for regional (A - Vancouver Island) and global pairs (B - worldwide)

**A**

| Pair | Habitat | Coordinates | Collection Date (dd/mm/yy) |
|---|---|---|---|
| Misty | Lake | 50° 36’ 20” N<br>127° 16’ 9” W | 23/5/13 |
| Misty | Stream | 50° 36’ 8” N<br>127° 15’ 1” W | 24/5/13 |
| Robert’s | Lake | 50° 12’ 58” N<br>125° 32’ 40” W | 26/5/13 |
| Robert’s | Stream | 50° 14’ 2” N<br>125° 34’ 7” W | 27/5/13 |
| Boot | Lake | 50° 3’ 38” N<br>125° 32’ 16” W | 29/5/13 |
| Boot | Stream | 50° 2’ 38” N<br>125° 33’ 39” W | 30/5/13 |
| Beaver | Lake | 50° 36’ 3” N<br>127° 18’ 37” W | 21/5/13 |
| Beaver | Stream | 50° 35’ 34” N<br>127° 21’ 2” W | 22/5/13 |
| Pye | Lake | 50° 17’ 21” N<br>125° 34’ 14” W | 25/5/13, 28/6/13 |
| Pye | Stream | 50° 18’ 50” N<br>125° 33’ 48” W | 27/5/13 |
| Village Bay | Lake | 50° 10’ 30” N<br>125° 11’ 25” W | 15/6/13, 27/6/13 |
| Village Bay | Stream | 50° 10’ 18” N<br>125° 11’ 52” W | 14/6/13 |

**B**

| Site | Habitat | Country | Locale | Coordinates | Date |
|---|---|---|---|---|---|
| Lake Ainslie | Lake | Canada | Cape Breton,<br>Nova Scotia | 46° 10’ 46” N<br>61° 16’ 17” W | 1/7/14 |
| Black River | Stream | Canada | Cape Breton,<br>Nova Scotia | 46° 9’ 28” N<br>61° 16’ 32” W | 6/6/14 |
| Lake Constance | Lake | Switzerland | Romanshorn,<br>Thurgau | 47° 33’ 22” N<br>9° 22’ 44” E | 21/5/14 |
| Aach River | Stream | Switzerland | Erlen,<br>Thurgau | 47° 32’ 54” N<br>9° 12’ 35” E | 21/5/14 |
| Lake Plon | Lake | Germany | Plon,<br>Schleswig-Holstein | 54° 8’ 49” N<br>10° 24’ 30” E | 15/4/1 |
| Karperbek | Stream | Germany | Plon,<br>Schleswig-Holstein | 54° 9’ 52” N<br>10° 20’ 37” E | 10/4/14 |
| Skogseidsvatnet | Lake | Norway | Fusa,<br>Hordaland | 60° 14’ 62” N<br>5° 54’ 72” E | 8/8/14 |
| Orra | Stream | Norway | Fusa,<br>Hordaland | 60° 15’ 34” N<br>5° 55’ 58” E | 8/8/14 |
| Flodid | Lake | Iceland | A-Húnavatnssýsla<br>Norðurland- vestra | 65° 29’ 21” N<br>20° 21’ 30” W | 25/7/14 |
| Vatnsdalsa | Stream | Iceland | A - Húnavatnssýsla<br>Norðurland- vestra | 65° 30’ 25” N<br>65° 29’ 21” W | 25/7/14 |
| Big Lake | Lake | USA | Matanuska-Susitna,<br>Alaska | 61° 32’ 3” N<br>149° 50’ 1” W | 16/6/14 |
| Fish Creek | Stream | USA | Matanuska-Susitna,<br>Alaska | 61° 32’ 2” N<br>149° 49’ 38” W | 16/6/14 |

**Table S2**. Description of linear morphological measurements

| Trait | Photograph | Description |
|---|---|---|
| First dorsal spine length | Lateral | Length, anterior insertion of spine to tip |
| Second dorsal spine length | Lateral | Length, anterior insertion of spine to tip |
| Dorsal fin length | Lateral | Length, at insertion points of fin |
| Caudal peduncle depth | Lateral | Depth, narrowest point of caudal peduncle |
| Anal fin length | Lateral | Length, at insertion points of fin |
| Pectoral fin length | Lateral | Length, dorsal to ventral insertion of fin |
| Body depth | Lateral | Depth, anterior insertion of first dorsal spine to anterior point of pelvic girdle |
| Mouth length | Lateral | Length, inside corner of mouth to anterior tip of lower jaw |
| Snout length | Lateral | Length, tip of upper jaw to closest point on the perimeter of the eye |
| Eye length | Lateral | Diameter, from posterior end of snout |
| Head length | Lateral | Length, tip of upper jaw to posterior-most point of the gill operculum |
| Pectoral fin width | Lateral | Width, measured from distal ends of outer fin rays |
| Pectoral fin length | Lateral | Length, longest fin ray |
| Standard length | Lateral | Length, tip of snout to end of caudal peduncle |
| Buccal cavity length | Ventral | Length, tip of snout to anterior point of ectocoracoid |
| Gape width | Ventral | Width, distance between mouth corners |
| Body width 1 | Ventral | Width, jawbone to jawbone, centered at the eyes |
| Body width 2 | Ventral | Width, distance between posterior points of ectocoracoid |
| Pelvic girdle width | Ventral | Width, between insertions of pelvic spines |
| Pelvic girdle length | Ventral | Length, anterior-most point of girdle to caudal tip of posterior process |
| Pelvic girdle width, posterior process | Ventral | Width, widest points of posterior process |
| Pelvic girdle length, posterior process | Ventral | Length, from anterior-most point to caudal tip |
| Body width 3 | Ventral | Width, across anterior insertion of anal fin |
| Body width 4 | Ventral | Width, across posterior insertion of anal fin |

**Table S3**. Results from MANCOVAs on the multivariate traits of regional (A- Vancouver Island) and global pairs (B). Fixed effects were habitat (Hab.), pair (Pair), sex (S), and centroid size (S), along with all two-way interactions. Degrees of freedom (df), Wilks' lambda (Wilks' λ), F values (*F*), and effect sizes (Partial $\eta^2$) are depicted. Significant F values ($P \leq 0.05$) are in bold.

**A**

**Univariate**

| | Hab. | Pair | S | C | Hab. x Pair | Hab. x S | Hab. x C | Pair x S | Pair x C | S x C | Error |
|---|---|---|---|---|---|---|---|---|---|---|---|
| df | 1 | 5 | 2 | 1 | 5 | 2 | 1 | 10 | 5 | 2 | 395 |
| Wilks' λ | 0.14 | 0.00 | 0.05 | 0.01 | 0.04 | 0.75 | 0.74 | 0.43 | 0.43 | 0.77 | |
| *F* | **85** | **43** | **47** | **1291** | **12** | **2** | **5** | 1 | **3** | **2** | |
| Partial $\eta^2$ | 0.81 | 0.55 | 0.39 | 0.98 | 0.35 | 0.11 | 0.24 | 0.08 | 0.15 | 0.12 | |

**Gill rakers**

| | Hab. | Pair | S | C | Hab. x Pair | Hab. x S | Hab. x C | Pair x S | Pair x C | S x C | Error |
|---|---|---|---|---|---|---|---|---|---|---|---|
| df | 1 | 5 | 2 | 1 | 5 | 2 | 1 | 10 | 5 | 2 | 444 |
| Wilks' λ | 0.38 | 0.52 | 0.85 | 0.94 | 0.48 | 0.96 | 0.99 | 0.84 | 0.93 | 0.97 | |
| *F* | **139** | **13** | **8** | 6 | **14** | 2 | 1 | 2 | 1 | 1 | |
| Partial $\eta^2$ | 0.45 | 0.13 | 0.09 | 0.02 | 0.08 | 0.03 | 0.01 | 0.04 | 0.02 | 0.02 | |

**Body shape**

| | Hab. | Pair | S | C | Hab. x Pair | Hab. x S | Hab. x C | Pair x S | Pair x C | S x C | Error |
|---|---|---|---|---|---|---|---|---|---|---|---|
| df | 1 | 5 | 2 | 1 | 5 | 2 | 10 | 1 | 5 | 2 | 444 |
| Wilks' λ | 0.11 | 0.02 | 0.27 | 0.31 | 0.04 | 0.76 | 0.38 | 0.82 | 0.56 | 0.82 | |
| *F* | **99.30** | **16.80** | **11.60** | **28.40** | **11.60** | **1.90** | **1.30** | **2.90** | **1.50** | **1.30** | |
| Partial $\eta^2$ | 0.85 | 0.52 | 0.40 | 0.64 | 0.37 | 0.12 | 0.08 | 0.16 | 0.11 | 0.09 | |

**B**

**Univariate**

| | Hab. | Pair | S | C | Hab. x Pair | Hab. x S | Hab. x C | Pair x S | Pair x C | S x C | Error |
|---|---|---|---|---|---|---|---|---|---|---|---|
| df | 1 | 5 | 1 | 1 | 5 | 1 | 1 | 5 | 5 | 1 | 453 |
| Wilks' $\lambda$ | 0.11 | 0.00 | 0.16 | 0.03 | 0.12 | 0.89 | 0.88 | 0.39 | 0.53 | 0.92 | |
| *F* | **127** | **93** | **83** | **541** | **9** | **2** | **2** | **3** | **2** | **1** | |
| Partial $\eta^2$ | 0.41 | 0.71 | 0.67 | 0.95 | 0.23 | 0.1 | 0.09 | 0.13 | 0.12 | 0.08 | |

**Gill rakers**

| | Hab. | Pair | S | C | Hab. x Pair | Hab. x S | Hab. x C | Pair x S | Pair x C | S x C | Error |
|---|---|---|---|---|---|---|---|---|---|---|---|
| df | 1 | 5 | 1 | 1 | 5 | 1 | 1 | 5 | 5 | 1 | 444 |
| Wilks' $\lambda$ | 0.59 | 0.11 | 0.93 | 0.37 | 0.69 | 0.98 | 0.99 | 0.93 | 0.86 | 0.99 | |
| *F* | **60** | **55** | **7** | **152** | **7** | 1 | 1 | 1 | **3** | 1 | |
| Partial $\eta^2$ | 0.031 | 0.11 | 0.115 | 0.452 | 0.033 | 0.008 | 0.008 | 0.016 | 0.028 | 0.007 | |

**Body shape**

| | Hab. | Pair | S | C | Hab. x Pair | Hab. x S | Hab. x C | Pair x S | Pair x C | S x C | Error |
|---|---|---|---|---|---|---|---|---|---|---|---|
| df | 1 | 5 | 1 | 1 | 5 | 1 | 5 | 1 | 5 | 1 | 453 |
| Wilks' $\lambda$ | 0.53 | 0.01 | 0.24 | 0.59 | 0.19 | 0.86 | 0.38 | 0.85 | 0.57 | 0.88 | |
| *F* | **11.52** | **29.85** | **41.26** | **8.64** | **4.97** | **2.15** | **2.67** | **2.19** | **1.51** | **1.66** | |
| Partial $\eta^2$ | 0.34 | 0.67 | 0.66 | 0.23 | 0.18 | 0.11 | 0.14 | 0.14 | 0.11 | 0.12 | |

**Table S4**. Results from ANCOVAs on single traits of regional (A- Vancouver Island) and global pairs (B). Fixed effects were habitat (Hab.), pair (Pair), sex (S), and centroid size (C), along with all two-way interactions. Degrees of freedom (df), F values (*F*), and effect sizes (Partial $\eta^2$) are depicted. Significant F values ($P \leq 0.05$) are in bold.

**A**

**Body depth**

| | Hab. | Pair | S | C | Hab. x Pair | Hab. x S | Hab. x C | Pair x S | Pair x C | S x C | Error |
|---|---|---|---|---|---|---|---|---|---|---|---|
| df | 1 | 5 | 2 | 1 | 5 | 2 | 1 | 10 | 5 | 2 | 444 |
| *F* | **1040.20** | **30.60** | **7.20** | **41.40** | **6.80** | 0.70 | **8.50** | 0.90 | 1.4 | 3.50 | |
| Partial $\eta^2$ | 0.48 | 0.12 | 0.01 | 0.02 | 0.02 | 0.00 | 0.01 | 0.01 | 0.01 | 0.01 | |

**Gill raker number**

| | Hab. | Pair | S | C | Hab. x Pair | Hab. x S | Hab. x C | Pair x S | Pair x C | S x C | Error |
|---|---|---|---|---|---|---|---|---|---|---|---|
| df | 1 | 5 | 2 | 1 | 5 | 2 | 1 | 10 | 5 | 2 | 444 |
| *F* | **552.40** | **24.60** | 1.60 | **10.40** | **26.00** | **3.80** | 1.70 | 0.80 | 1.00 | 0.50 | |
| Partial $\eta^2$ | 0.29 | 0.13 | 0.00 | 0.01 | 0.09 | 0.01 | 0.00 | 0.01 | 0.01 | 0.00 | |

**Mean gill raker length**

| | Hab. | Pair | S | C | Hab. x Pair | Hab. x S | Hab. x C | Pair x S | Pair x C | S x C | Error |
|---|---|---|---|---|---|---|---|---|---|---|---|
| df | 1 | 5 | 2 | 1 | 5 | 2 | 1 | 10 | 5 | 2 | 444 |
| *F* | **32.50** | **29.70** | **31.70** | **8.60** | **37.70** | 1.20 | 0.10 | **4.90** | **4.00** | 2.20 | |
| Partial $\eta^2$ | 0.02 | 0.17 | 0.10 | 0.00 | 0.10 | 0.00 | 0.00 | 0.06 | 0.02 | 0.01 | |

**Gill raker spacing**

| | Hab. | Pair | S | C | Hab. x Pair | Hab. x S | Hab. x C | Pair x S | Pair x C | S x C | Error |
|---|---|---|---|---|---|---|---|---|---|---|---|
| df | 1 | 5 | 2 | 1 | 5 | 2 | 1 | 10 | 5 | 2 | 444 |
| *F* | **132.20** | **12.80** | **8.00** | **3.60** | **17.00** | 1.10 | 0.10 | **1.90** | 0.80 | 1.40 | |

| | | | | | | | | | | |
|---|---|---|---|---|---|---|---|---|---|---|
| Partial $\eta^2$ | 0.10 | 0.10 | 0.02 | 0.00 | 0.08 | 0.01 | 0.00 | 0.02 | 0.01 | 0.00 |

**B**

**Body depth**

| | Hab. | Pair | S | C | Hab. x Pair | Hab. x S | Hab. x C | Pair x S | Pair x C | S x C | Error |
|---|---|---|---|---|---|---|---|---|---|---|---|
| df | 1 | 5 | 1 | 1 | 5 | 1 | 1 | 5 | 5 | 1 | 453 |
| *F* | **90.00** | **246.90** | 2.90 | **14.10** | **14.30** | 0.20 | **7.10** | 1.20 | 0.30 | 0.60 | |
| Partial $\eta^2$ | 0.02 | 0.67 | 0.00 | 0.00 | 0.03 | 0.00 | 0.00 | 0.00 | 0.00 | 0.00 | |

**Gill raker number**

| | Hab. | Pair | S | C | Hab. x Pair | Hab. x S | Hab. x C | Pair x S | Pair x C | S x C | Error |
|---|---|---|---|---|---|---|---|---|---|---|---|
| df | 1 | 5 | 1 | 1 | 5 | 1 | 1 | 5 | 5 | 1 | 453 |
| *F* | **30.20** | **59.70** | 4.50 | **5.70** | **7.00** | 0.00 | 0.20 | 1.50 | 1.40 | 0.90 | |
| Partial $\eta^2$ | 0.01 | 0.21 | 0.01 | 0.00 | 0.03 | 0.00 | 0.00 | 0.01 | 0.01 | 0.00 | |

**Mean Gill raker length**

| | Hab. | Pair | S | C | Hab. x Pair | Hab. x S | Hab. x C | Pair x S | Pair x C | S x C | Error |
|---|---|---|---|---|---|---|---|---|---|---|---|
| df | 1 | 5 | 1 | 1 | 5 | 1 | 1 | 5 | 5 | 1 | 444 |
| *F* | **142.50** | **179.10** | **19.80** | **319.20** | **9.70** | 0.00 | 0.70 | **1.90** | 1.10 | 0.30 | |
| Partial $\eta^2$ | 0.00 | 0.12 | 0.07 | 0.15 | 0.04 | 0.00 | 0.00 | 0.01 | 0.01 | 0.00 | |

**Gill raker spacing**

| | Hab. | Pair | S | C | Hab. x Pair | Hab. x S | Hab. x C | Pair x S | Pair x C | S x C | Error |
|---|---|---|---|---|---|---|---|---|---|---|---|
| df | 1 | 5 | 1 | 1 | 5 | 1 | 1 | 5 | 5 | 1 | 444 |
| *F* | **115.00** | **257.60** | 5.80 | **518.00** | **8.10** | 2.50 | 2.30 | 1.20 | **9.20** | 0.80 | |
| Partial $\eta^2$ | 0.00 | 0.06 | 0.01 | 0.33 | 0.02 | 0.00 | 0.00 | 0.01 | 0.05 | 0.00 | |

**Table S5**. Mean pairwise $F_{ST}$ values across pairs at the regional (A) and global (B) scales. Abbreviations stand for Village Bay (Bay), Beaver (Beav), Boot (Boot), Misty (Mist), Pye (Pye), Robert (Rob), Nova Scotia (Nos), Germany (Ger), Iceland (Ice), Norway (Nor), Switzerland (Swis), and Alaska (AK). Lake and stream populations are represented by (-L) and (-S), respectively.

**A**

| | Bay-S | Beav-L | Beav-S | Boot-L | Boot-S | Mist-L | Mist-S | Pye-L | Pye-S | Rob-L | Rob-S |
|---|---|---|---|---|---|---|---|---|---|---|---|
| Bay-L | 0.026 | 0.039 | 0.050 | 0.074 | 0.063 | 0.080 | 0.075 | 0.039 | 0.039 | 0.084 | 0.081 |
| Bay-S | | 0.053 | 0.062 | 0.089 | 0.078 | 0.097 | 0.091 | 0.056 | 0.056 | 0.099 | 0.097 |
| Beav-L | | | 0.025 | 0.068 | 0.059 | 0.083 | 0.078 | 0.044 | 0.044 | 0.090 | 0.086 |
| Beav-S | | | | 0.081 | 0.070 | 0.098 | 0.092 | 0.059 | 0.058 | 0.103 | 0.100 |
| Boot-L | | | | | 0.055 | 0.147 | 0.135 | 0.086 | 0.086 | 0.147 | 0.147 |
| Boot-S | | | | | | 0.134 | 0.122 | 0.074 | 0.073 | 0.135 | 0.134 |
| Mist-L | | | | | | | 0.007 | 0.073 | 0.079 | 0.050 | 0.045 |
| Mist-S | | | | | | | | 0.066 | 0.071 | 0.046 | 0.042 |
| Pye-L | | | | | | | | | 0.008 | 0.081 | 0.077 |
| Pye-S | | | | | | | | | | 0.085 | 0.081 |
| Rob-L | | | | | | | | | | | 0.014 |

**B**

| | Nos-S | Ger-L | Ger-S | Ice-L | Ice-S | Nor-L | Nor-S | Swis-L | Swis-S | AK-L | AK-S |
|---|---|---|---|---|---|---|---|---|---|---|---|
| Nos-L | 0.037 | 0.030 | 0.046 | 0.055 | 0.061 | 0.094 | 0.091 | 0.093 | 0.067 | 0.123 | 0.130 |
| Nos-S | | 0.035 | 0.041 | 0.061 | 0.064 | 0.110 | 0.107 | 0.107 | 0.082 | 0.134 | 0.138 |
| Ger-L | | | 0.023 | 0.044 | 0.050 | 0.082 | 0.080 | 0.084 | 0.061 | 0.120 | 0.130 |
| Ger-S | | | | 0.058 | 0.061 | 0.103 | 0.100 | 0.104 | 0.080 | 0.135 | 0.142 |
| Ice-L | | | | | 0.029 | 0.101 | 0.099 | 0.100 | 0.079 | 0.130 | 0.141 |
| Ice-S | | | | | | 0.110 | 0.109 | 0.108 | 0.088 | 0.136 | 0.144 |
| Nor-L | | | | | | | 0.011 | 0.061 | 0.041 | 0.079 | 0.136 |
| Nor-S | | | | | | | | 0.057 | 0.037 | 0.080 | 0.133 |
| Swis-L | | | | | | | | | 0.032 | 0.081 | 0.124 |
| Swis-S | | | | | | | | | | 0.084 | 0.126 |
| AK-L | | | | | | | | | | | 0.109 |

**Table S6**. Pairwise observed $\theta_P$ (upper triangle) and $\theta_G$ (lower triangle) at the regional (A) and global (B) scales. Abbreviations stand for Nova Scotia (Nos), Germany (Ger), Iceland (Ice), Norway (Nor), Switzerland (Swis), Alaska (AK), Village Bay (Bay), Beaver (Beav), Boot (Boot), Misty (Mist), Pye (Pye), and Robert (Rob).

**A**

| | Rob | Mist | Beav | Pye | Bay | Boot |
|---|---|---|---|---|---|---|
| Rob | | 65.03 | 46.07 | 45.95 | 63.60 | 83.94 |
| Mist | 91.51 | | 59.53 | 70.36 | 50.65 | 57.87 |
| Beav | 88.01 | 87.32 | | 52.77 | 58.44 | 79.41 |
| Pye | 87.43 | 87.49 | 92.36 | | 57.07 | 81.07 |
| Bay | 87.03 | 86.23 | 93.16 | 93.27 | | 52.77 |
| Boot | 91.84 | 92.59 | 87.61 | 87.15 | 86.75 | |

**B**

| | Ger | Swis | AK | Nos | Ice | Nor |
|---|---|---|---|---|---|---|
| Ger | | 65.03 | 106.63 | 93.51 | 102.16 | 68.30 |
| Swis | 91.85 | | 97.35 | 95.80 | 73.40 | 68.47 |
| AK | 91.04 | 90.47 | | 112.19 | 82.05 | 92.82 |
| Nos | 93.05 | 92.31 | 91.44 | | 85.37 | 114.65 |
| Ice | 87.21 | 87.15 | 88.24 | 88.18 | | 94.94 |
| Nor | 89.33 | 90.02 | 89.67 | 91.04 | 89.49 | |

**Table S7**. Pairwise observed $\Delta L_P$ (upper triangle) and $\Delta L_G$ (lower triangle) at the regional (A) and global (B) scales. Abbreviations stand for Nova Scotia (Nos), Germany (Ger), Iceland (Ice), Norway (Nor), Switzerland (Swis), Alaska (AK), Village Bay (Bay), Beaver (Beav), Boot (Boot), Misty (Mist), Pye (Pye), and Robert (Rob).

**A**

| | Rob | Mist | Beav | Pye | Bay | Boot |
|---|---|---|---|---|---|---|
| Rob | | 4.58 | -8.76 | -16.12 | -0.29 | -6.09 |
| Mist | -0.69 | | -13.34 | 20.70 | 4.29 | -10.68 |
| Beav | -2.41 | -1.72 | | 7.36 | -9.05 | -2.67 |
| Pye | -1.11 | -0.42 | 1.29 | | -16.41 | 10.03 |
| Bay | -0.19 | 0.49 | 2.21 | 0.92 | | -6.38 |
| Boot | 0.82 | 1.51 | 3.2 | 1.93 | 1.01 | |

**B**

| | Ger | Swis | AK | Nos | Ice | Nor |
|---|---|---|---|---|---|---|
| Ger | | 10.01 | 4.34 | 6.89 | -9.06 | 2.63 |
| Swis | -0.19 | | -5.67 | 16.90 | 19.07 | 7.38 |
| AK | 0.02 | 0.22 | | 11.23 | 13.40 | 1.70 |
| Nos | 0.97 | 1.17 | 0.95 | | -2.17 | 9.52 |
| Ice | 3.41 | 3.61 | 3.38 | 2.44 | | 11.69 |
| Nor | 16.01 | 16.19 | 15.98 | 15.03 | 12.59 | |

**Table S8**. Pairwise observed $\theta_{OUTLIERS}$ at the regional (A) and global (B) scales. Abbreviations stand for Nova Scotia (Nos), Germany (Ger), Iceland (Ice), Norway (Nor), Switzerland (Swis), Alaska (AK), Village Bay (Bay), Beaver (Beav), Boot (Boot), Misty (Mist), Pye (Pye), and Robert (Rob).

**A**

| | Rob | Mist | Beav | Pye | Bay | Boot |
|---|---|---|---|---|---|---|
| Rob | | | | | | |
| Mist | 85.66 | | | | | |
| Beav | 97.35 | 85.72 | | | | |
| Pye | 84.23 | 84.28 | 84.79 | | | |
| Bay | 86.41 | 84.91 | 84.92 | 71.05 | | |
| Boot | 92.07 | 87.43 | 91.38 | 80.44 | 82.05 | |

**B**

| | Ger | Swis | AK | Nos | Ice | Nor |
|---|---|---|---|---|---|---|
| Ger | | | | | | |
| Swis | 88.81 | | | | | |
| AK | 86.01 | 92.19 | | | | |
| Nos | 93.79 | 87.21 | 87.37 | | | |
| Ice | 89.89 | 90.24 | 89.55 | 92.02 | | |
| Nor | 86.23 | 95.86 | 79.69 | 90.64 | 87.77 | |

**Table S9**. Pairwise observed $\Delta L_{OUTLIERS}$ at the regional (A) and global (B) scales. Abbreviations stand for Nova Scotia (Nos), Germany (Ger), Iceland (Ice), Norway (Nor), Switzerland (Swis), Alaska (AK), Village Bay (Bay), Beaver (Beav), Boot (Boot), Misty (Mist), Pye (Pye), and Robert (Rob).

**A**

| | Rob | Mist | Beav | Pye | Bay | Boot |
|---|---|---|---|---|---|---|
| Rob | | | | | | |
| Mist | 3.67 | | | | | |
| Beav | 7.57 | 3.91 | | | | |
| Pye | 7.93 | 4.26 | 0.36 | | | |
| Bay | 2.83 | -0.84 | -4.74 | -5.11 | | |
| Boot | 4.92 | 1.25 | -2.65 | -3.01 | 2.09 | |

**B**

| | Ger | Swis | AK | Nos | Ice | Nor |
|---|---|---|---|---|---|---|
| Ger | | | | | | |
| Swis | 2.41 | | | | | |
| AK | 2.64 | 0.24 | | | | |
| Nos | 0.91 | -1.49 | -1.74 | | | |
| Ice | 4.42 | 2.02 | 1.78 | 3.52 | | |
| Nor | 30.01 | 27.61 | 27.37 | 29.11 | 25.59 | |

### Figure legends

**Figure S1**: Maps of study sites for regional, Vancouver Island (A) and worldwide, global samples (B).

**Figure S2**: Locations of the 19 landmarks for geometric morphometrics.

**Figure S3**: Standard deviations (st.dev.) of lake and stream environmental variables calculated within watersheds at both global (black) and regional (grey) scales. Significant differences are indicated with two stars when $P < 0.001$ and one star when $P < 0.05$. Actual values are depicted above each bar.

**Figure S4**: Among watershed variation in lake and stream environmental variables at both global (black) and regional (grey) scales. Actual values are depicted above each bar.

**Figure S5**: Neighbor-joining trees of regional (A) and global (B) lake-stream pairs based on average pairwise $F_{ST}$ values of none-outlier SNPs.

**Figure S1**

**A**

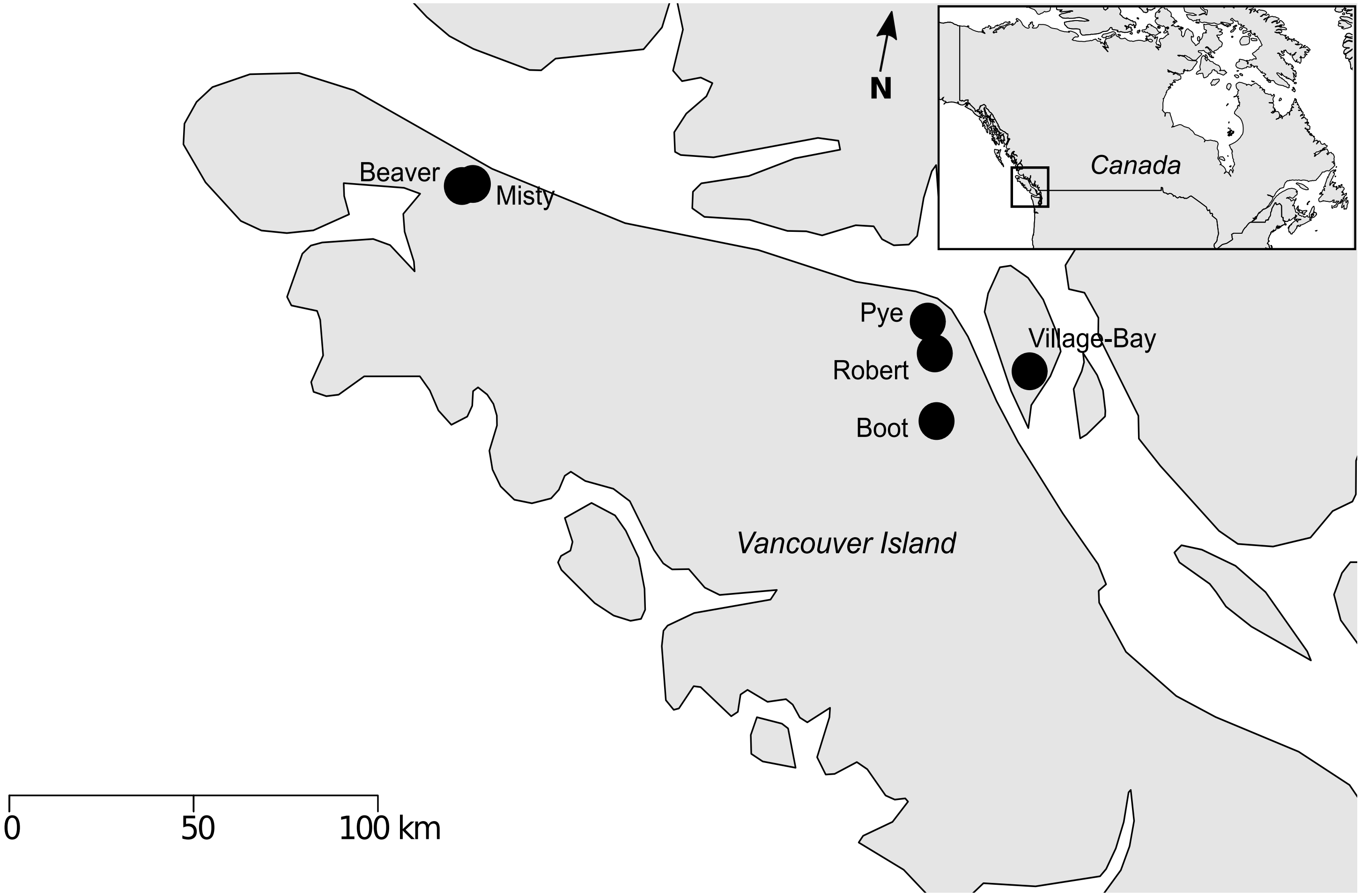
N
Canada
Beaver
Misty
Pye
Robert
Boot
Village-Bay
Vancouver Island
0
50
100 km

**B**

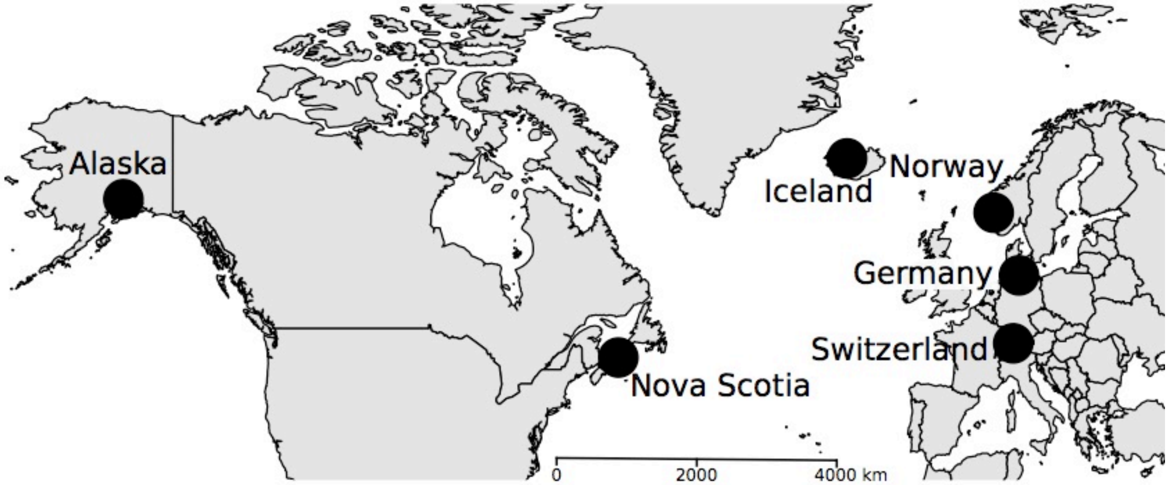
Alaska
Iceland
Norway
Germany
Switzerland
Nova Scotia
0
2000
4000 km

**Figure S2**

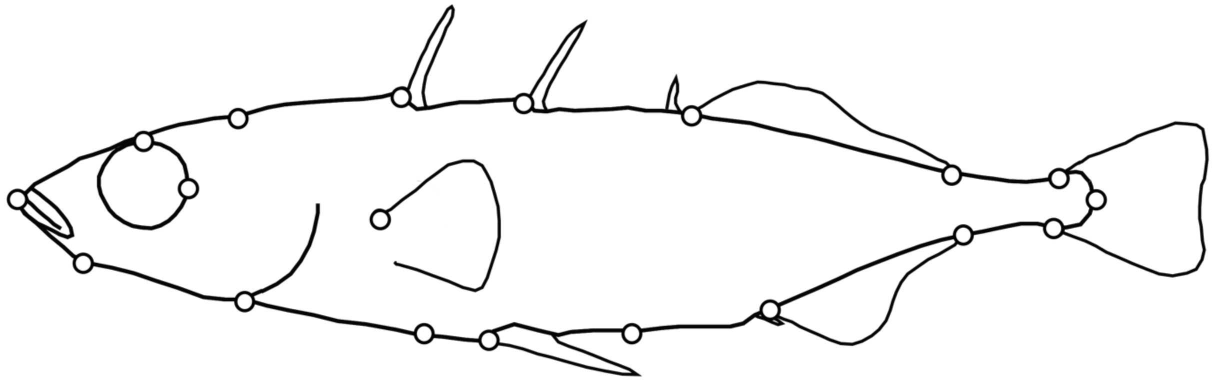

**Figure S3**

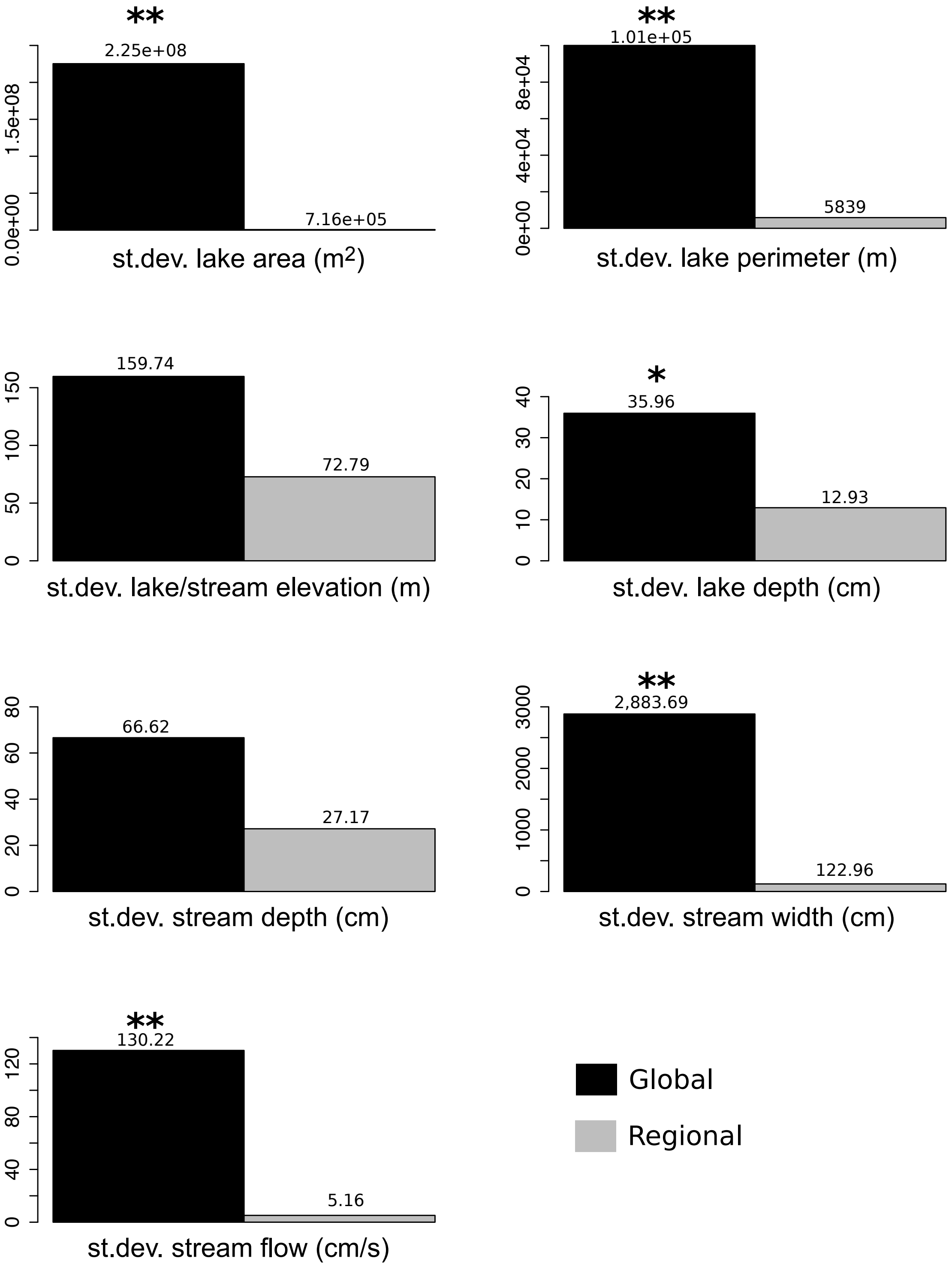
**
2.25e+08
7.16e+05
0.0e+00
1.5e+08
st.dev. lake area (m2)
**
1.01e+05
5839
0e+00
4e+04
8e+04
st.dev. lake perimeter (m)
159.74
72.79
0
50
100
150
st.dev. lake/stream elevation (m)
*
35.96
12.93
0
10
20
30
40
st.dev. lake depth (cm)
66.62
27.17
0
20
40
60
80
st.dev. stream depth (cm)
**
2,883.69
122.96
0
1000
2000
3000
st.dev. stream width (cm)
**
130.22
5.16
0
40
80
120
st.dev. stream flow (cm/s)
Global
Regional

**Figure S4**

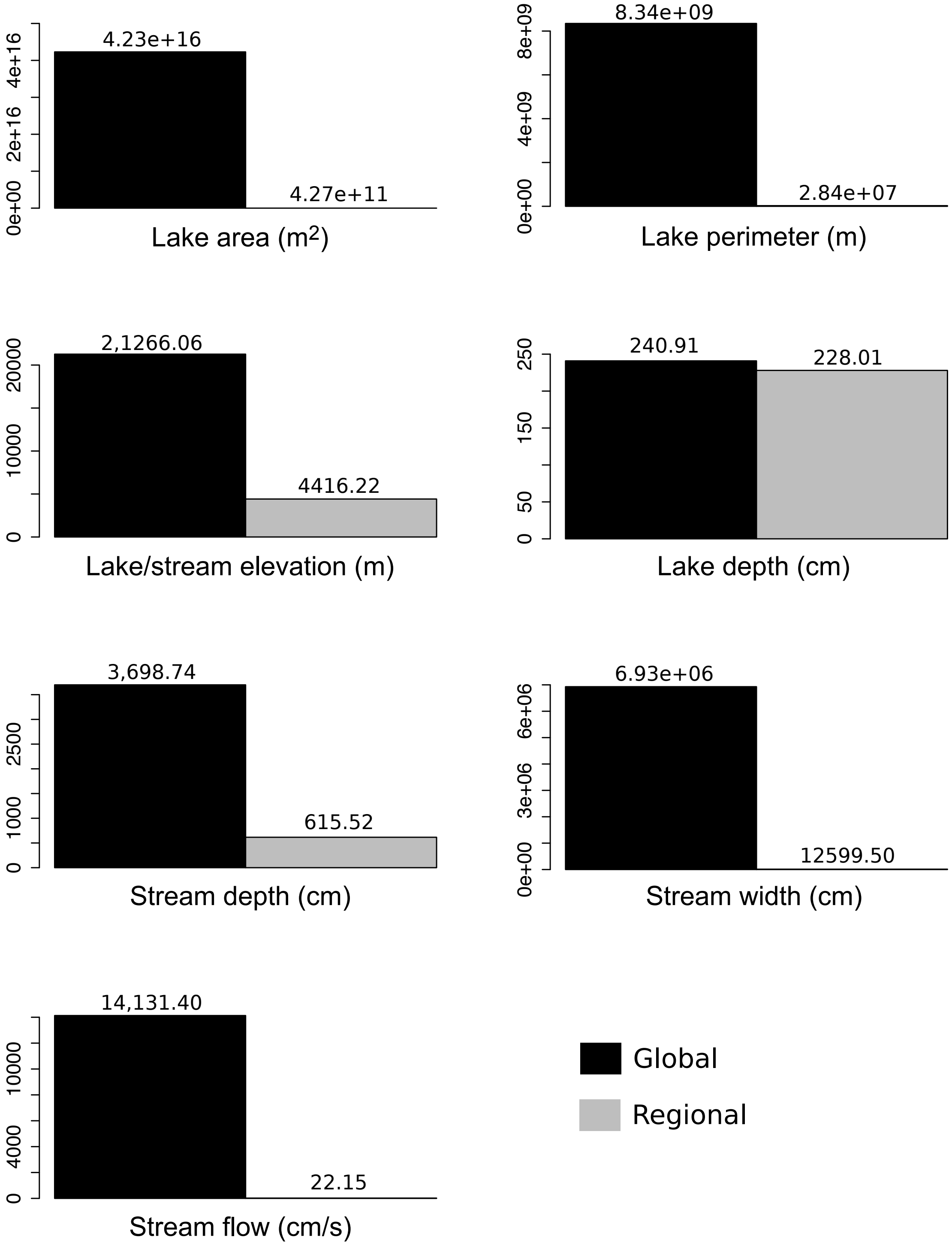
4.23e+16
4.27e+11
0e+00 2e+16 4e+16
Lake area (m2)
8.34e+09
2.84e+07
0e+00 4e+09 8e+09
Lake perimeter (m)
2,1266.06
4416.22
0 10000 20000
Lake/stream elevation (m)
240.91
228.01
0 50 150 250
Lake depth (cm)
3,698.74
615.52
0 1000 2500
Stream depth (cm)
6.93e+06
12599.50
0e+00 3e+06 6e+06
Stream width (cm)
14,131.40
22.15
0 4000 10000
Stream flow (cm/s)
Global
Regional

**Figure S5**

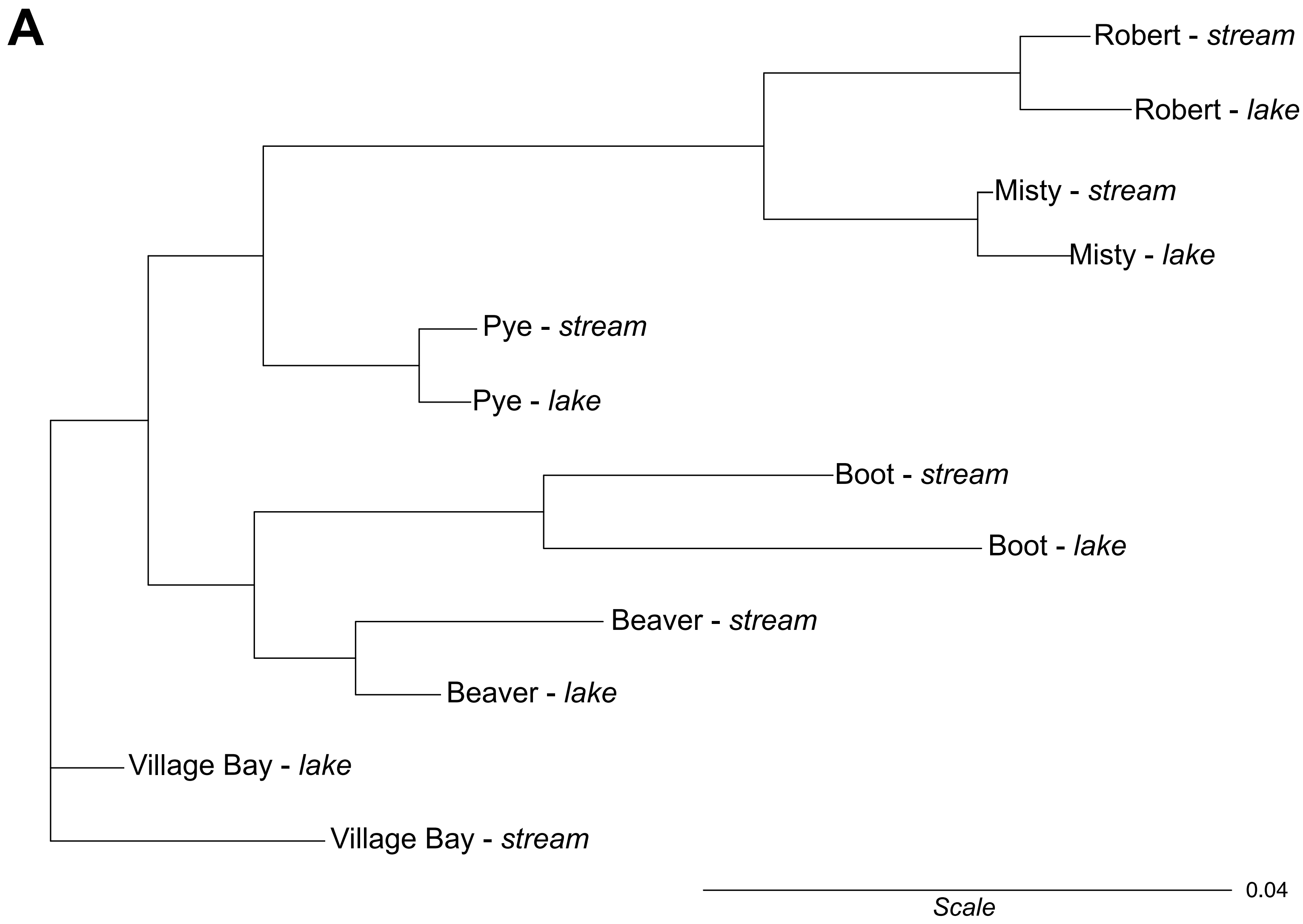
A
Robert - *stream*
Robert - *lake*
Misty - *stream*
Misty - *lake*
Pye - *stream*
Pye - *lake*
Boot - *stream*
Boot - *lake*
Beaver - *stream*
Beaver - *lake*
Village Bay - *lake*
Village Bay - *stream*
0.04
*Scale*

**B**

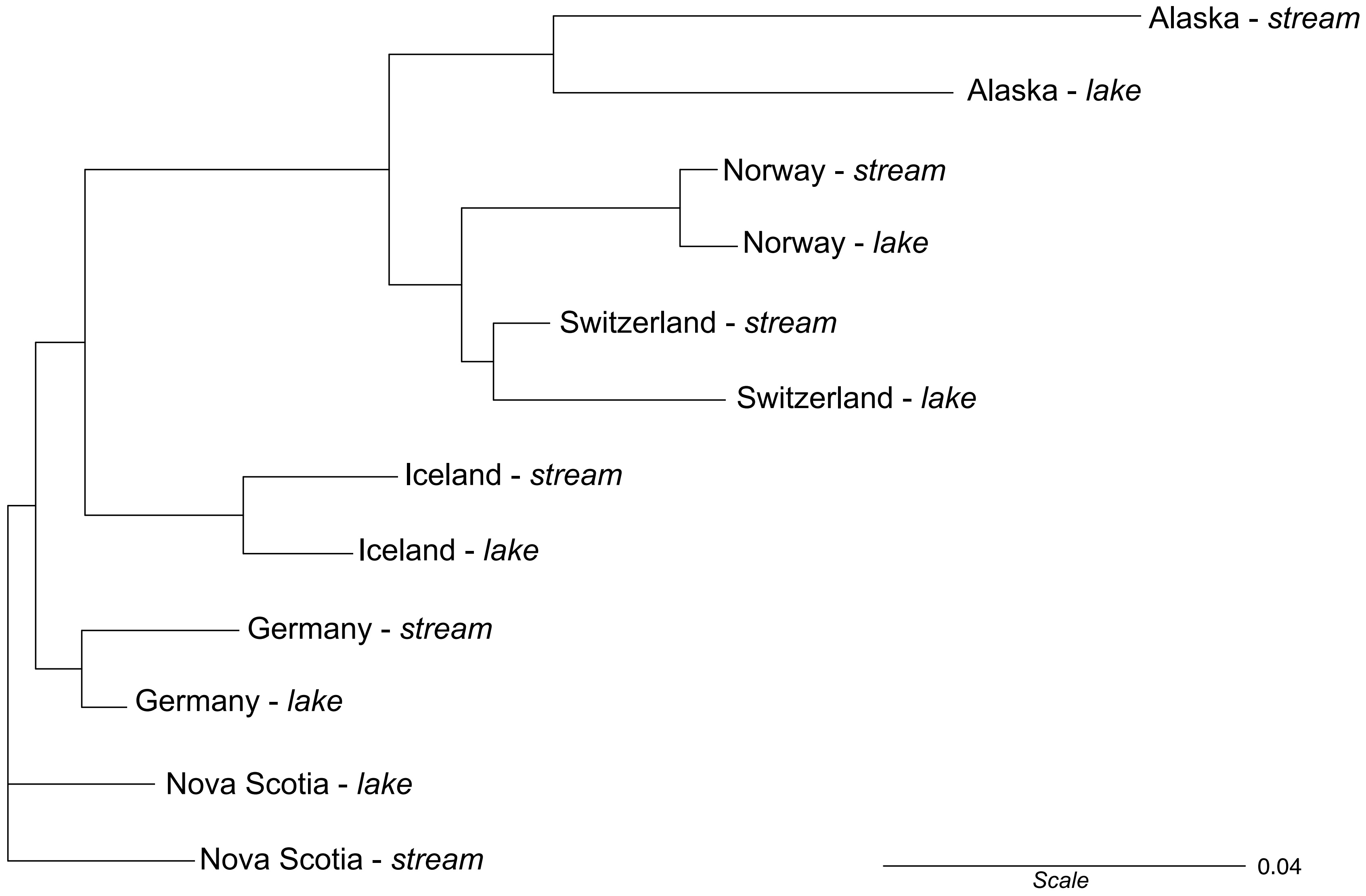
Alaska - *stream*
Alaska - *lake*
Norway - *stream*
Norway - *lake*
Switzerland - *stream*
Switzerland - *lake*
Iceland - *stream*
Iceland - *lake*
Germany - *stream*
Germany - *lake*
Nova Scotia - *lake*
Nova Scotia - *stream*
0.04
*Scale*
